# De novo pyrimidine biosynthesis inhibition synergizes with BCL-X_L_ targeting in pancreatic cancer

**DOI:** 10.1101/2024.05.17.594738

**Authors:** Huan Zhang, Qijia Yu, Naiara Santana-Codina, Clara Poupault, Claudia Campos, Xingping Qin, Aparna Padhye, Nicole Sindoni, Miljan Kuljanin, Junning Wang, Matthew J. Dorman, Andrew J. Aguirre, Stephanie K. Dougan, Kristopher A. Sarosiek, Joseph D. Mancias

## Abstract

Oncogenic KRAS, the genetic driver of 90% of pancreatic adenocarcinoma (PDAC), induces a metabolic rewiring characterized, in part, by dependency on de novo pyrimidine biosynthesis. Pharmacologic inhibition of dihydroorotate dehydrogenase (DHODH), an enzyme in the de novo pyrimidine synthesis pathway, delays pancreatic tumor growth in vivo; however, limited monotherapy efficacy suggests compensatory pathways and that combinatorial strategies are required for enhanced efficacy. Here, we use an integrated metabolomic, quantitative temporal proteomic and in vitro and in vivo DHODH inhibitor anchored CRISPR/Cas9 genetic screening approach to identify compensatory pathways to DHODH inhibition (DHODHi) and targets for combination strategies. We demonstrate that DHODHi alters the apoptotic regulatory proteome thereby enhancing sensitivity to inhibitors of the anti-apoptotic BCL2L1 (BCL-X_L_) protein. Combinatorial regimens with DHODH and BCL-X_L_ inhibition synergistically induce apoptosis in PDAC cell lines and patient-derived PDAC organoids. In vivo DHODH inhibition with Brequinar and BCL-X_L_ degradation with DT2216, a proteolysis targeting chimera (PROTAC), significantly inhibits the growth of PDAC tumors. Our data defines mechanisms of adaptation to DHODH inhibition and identifies a combination therapy strategy in PDAC.

## INTRODUCTION

Metabolic rewiring is a hallmark of cancer^1^. In pancreatic adenocarcinoma (PDAC), this metabolic shift is driven by oncogenic KRAS mutation that promotes glucose use for glycosylation and synthesis of nucleotides^2,3^. Nucleotide metabolism is critical for maintaining essential pathways that support tumor growth like DNA synthesis, DNA damage repair, and glycosylation^4^. Pyrimidine nucleotides (UTP, CTP, TTP) can be obtained through two main pathways: the de novo pyrimidine synthesis pathway, which requires precursors like glutamine, aspartate, glucose-derived metabolites, and bicarbonate to form the pyrimidine ring, and the salvage pathway, which can import or recycle nucleobases and nucleosides^5^. Drugs targeting nucleotide metabolism (e.g., gemcitabine, 5-fluorouracil) are cornerstones of the two most effective chemotherapy regimens for PDAC patients (e.g., FOLFIRINOX and gemcitabine with nab-paclitaxel); however, therapeutic resistance to these chemotherapies is a significant clinical challenge.

We and others have previously identified de novo pyrimidine nucleotide synthesis as an in vivo metabolic vulnerability in PDAC^2,3,6,7^, which suggested that blocking pyrimidine synthesis may be a promising therapeutic strategy for PDAC. Dihydroorotate dehydrogenase (DHODH) is an enzyme in the *de novo* pyrimidine biosynthesis pathway with clinically available inhibitors. DHODH is located in the inner mitochondrial membrane and couples oxidation of dihydroorotate to orotate, a critical intermediate in the synthesis of pyrimidines, with reduction of ubiquinone. In addition, pyrimidines are essential for Complex I activity by promoting PDH and tricarboxylic acid cycle activities^8^. Therefore, DHODH inhibitors (DHODHi) target pyrimidine synthesis and mitochondrial respiration, an additional essential PDAC metabolic pathway. Prior unsuccessful clinical trials evaluating de novo pyrimidine biosynthesis inhibition as an anticancer strategy likely failed due to inadequate dosing strategies that prevented consistent pyrimidine depletion^9–12^ and use of low-potency inhibitors like leflunomide (NCT02509052, NCT01611675). More recently there has been a renewed interest in repurposing Brequinar (BQ), an FDA-approved DHODH inhibitor with improved potency compared to leflunomide, as well as other newly available DHODHi as anticancer therapeutics. We previously demonstrated that DHODHi reduced pyrimidine nucleotides and decreased clonogenic ability of PDAC cells with a therapeutic index over non-tumor cells^3^. However, despite in vivo on-target depletion of pyrimidines, DHODHi demonstrated only partial responses in a xenograft PDAC model^3^ suggesting intrinsic or acquired therapeutic resistance.

Therapeutic resistance is one of the most difficult clinical problems in PDAC with a wide variety of genetic and non-genetic mechanisms reported^13^. Identifying and targeting non-genetic mechanisms of resistance is a challenge that requires a multidisciplinary approach encompassing identification of metabolic, transcriptomic and proteomic alterations that promote survival despite on-target inhibition^14,15^.

Here, we identify pathways of adaptation to DHODHi using an integrated MS-based quantitative temporal proteomics workflow, liquid chromatography (LC)/MS-based metabolomics and an in vitro and in vivo DHODHi-anchored CRISPR/Cas9 genetic screen approach (**Fig. 1a**). Proteomics and metabolomics confirm on-target inhibition of pyrimidine synthesis as well as an expected compensatory upregulation of the nucleotide salvage pathway. Using a genome-wide *in vitro* DHODHi-anchored screen, we identify sgRNAs targeting the anti-apoptotic gene *BCL2L1* (*BCL-X_L_*) as a high priority synthetic lethal combinatorial hit. To account for differences in the in vitro versus in vivo metabolic milieu and prioritize candidates for translation, we perform a DHODHi-anchored mini-pool in vivo PDAC xenograft CRISPR/Cas9 screen thereby validating BCL-X_L_ as a high priority target. We demonstrate synergistic cytotoxicity and tumor growth inhibition in PDAC cells, patient-derived organoids, and mouse models of PDAC when DHODHi are combined with a BCL-X_L_ inhibitor (A-1331852) or degrader (DT2216). Given the growing interest in targeting de novo pyrimidine synthesis as a pan-cancer dependency, we propose a combination strategy of DHODH and BCL-X_L_ inhibition that can induce anti-tumor responses with potential clinical translatability.

**Figure 1.**
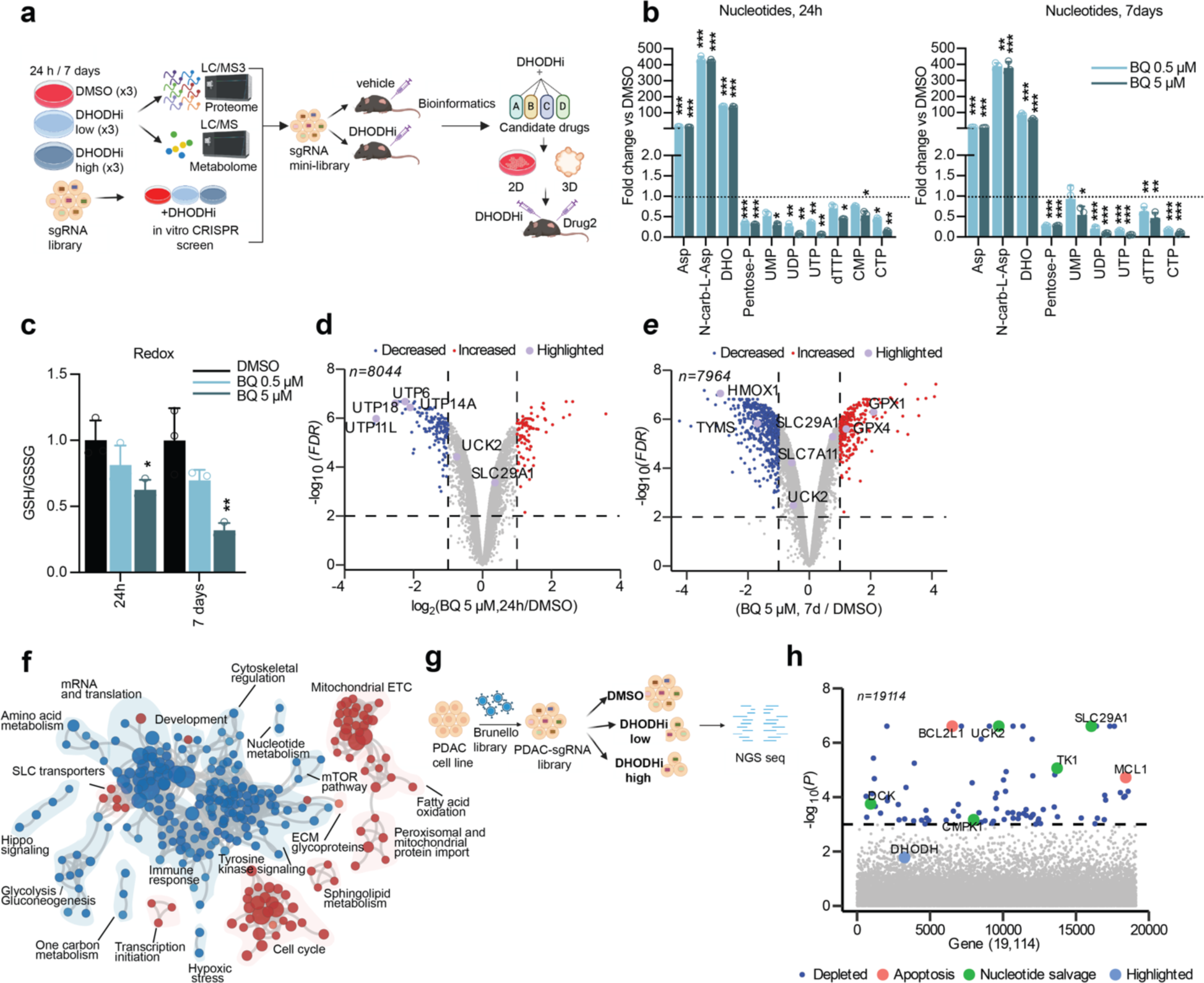
Defining mechanisms of short- and long-term adaptation to DHODH inhibition by an integrated multi-omics approach. **(a)** Schematic model of the multi-omic workflow used to define adaptation to DHODH inhibition using Brequinar (BQ). PDAC cells (PaTu-8988T and PaTu-8902) treated with low (0.5 μM) and high (5 μM) doses of BQ for 24h and 7 days were analyzed by mass spectrometry-based multiplexed tandem mass tag (TMT) quantitative proteomics and LC-MS/MS metabolomics. Synthetic lethalities were determined by a BQ-anchored genome scale loss-of-function CRISPR/Cas9 screen. High priority targets derived from the three techniques were used to build an sgRNA mini-library to evaluate in BQ-anchored in vivo xenograft experiments. Combinatorial strategies targeting candidate hits with available drugs were tested in vitro (2D and patient-derived organoid models) and in vivo in xenograft and orthotopic PDAC models. **(b-c)** Short and long-term BQ treatment demonstrates on-target metabolic effects. Values are represented as fold change of 24h and 7-day BQ-treated cells versus DMSO-treated cells at low (0.5 μM) and high dose (5 μM) BQ. Significant metabolites from each pathway (nucleotide and redox) are represented. Error bars represent s.d. of n=3 technical replicates from independently prepared samples from individual wells. Significance determined with t-test for cells treated with BQ vs. vehicle (*p < 0.05, **p < 0.01, ***p < 0.001). **(b)** Fold change of metabolites in the pyrimidine synthesis pathway correlate with a block in the pathway acutely (24h, left panel) and long-term (7 days, right panel). Asp: aspartate; N-Carb-L-Asp: N-Carbamoyl-L-Aspartate; DHO: dihydroorotate; UMP/UDP/UTP, uridine mono/di/triphosphate; CTP, cytidine triphosphate; dTTP, deoxythymidine triphosphate. **(c)** GSH/GSSG ratio is decreased at 24h and 7 days of BQ treatment. **(d-e)** Volcano plot illustrates significant protein abundance differences in PaTu-8988T cells treated with 5 μM BQ at 24h **(d)** and 7 days **(e)**. Volcano plots display the -log_10_ (FDR) versus the log_2_ of the relative protein abundance of mean BQ to DMSO-treated samples. Purple circles highlight proteins previously linked to DHODH function. Red circles represent significantly upregulated proteins (log_2_ fold change≥1, FDR<0.01), whereas blue circles represent significantly downregulated proteins (log_2_ fold change≤-1); data from 3 DMSO or 3 BQ-treated independent plates. **(f)** Enrichment map of gene set enrichment analysis (GSEA) of BQ-proteomes from PaTu-8988T cells at 7 days (0.5 μM). FDR<0.01, Jaccard coefficient>0.25, node size is related to the number of components identified within a gene set and the width of the line is proportional to the overlap between related gene sets. GSEA terms associated with upregulated (red), and downregulated (blue) proteins are colored accordingly and grouped into nodes with associated terms. **(g)** Schematic for BQ-anchored whole-genome in vitro screen. PaTu-8988T cells were lentivirally transduced with the Brunello library (19,114 genes, 76,441 sgRNAs), treated with BQ at 0.5 or 5 μM for 2 weeks and the remaining cells were analyzed by next-generation sequencing. **(h)** Manhattan plot for depleted hits in PaTu-8988T cells at 5 μM BQ. Blue dots highlight significant genes (-log_10_(P)=3) in BQ *vs*. DMSO, orange dots highlight genes related to apoptosis and green dots highlight genes in nucleotide salvage.

## RESULTS

### Defining adaptation to DHODHi using metabolomics and quantitative temporal proteomics

To define mechanisms of adaptation to DHODH inhibition, we initially tested sensitivity to Brequinar (BQ) in a panel of human and murine PDAC cell lines (**Supplementary Fig. 1a**). Half maximal inhibitory concentration (IC50) measurements showed a range of responses between 0.5-5 μM. To assess long-term responses that allow for analysis of adaptation at later time points, we analyzed doubling events in PaTu-8988T and PaTu-8902 cells treated with BQ for 12 days **(Supplementary Fig. 1b)**. Despite slower proliferation, PDAC cells maintained their proliferative capacity even at higher doses of BQ. Based on the varying levels of sensitivity in each cell line, we selected 0.5 and 5 μM as a low and high dose, respectively, for subsequent experiments.

First, we analyzed the metabolic response to BQ acutely (24 h) or long-term (7 days) by targeted LC-MS/MS metabolomics **(Supplementary Table 1)**. In agreement with previous studies^3^, BQ efficiently blocked pyrimidine synthesis (**Fig. 1b**) with accumulation of nucleotide precursors (Aspartate, N-Carbamoyl-Aspartate, Dihydroorotate) and a decrease in pyrimidine nucleotides (UMP, UDP, UTP, dTTP, CMP, CTP). Importantly, pyrimidine synthesis was consistently downregulated long-term, indicating adaptation to DHODHi was not due to loss of on-target DHODH inhibition and pathway reactivation. Long-term BQ increased metabolites in the nucleotide salvage pathway including uracil and uridine **(Supplementary Fig. 1c)**, suggesting compensatory activation of nucleoside import under pyrimidine depletion conditions^16^. DHODH catalyzes ubiquinone reduction and it has also been shown to support respiration by modulating PDH activity^3,8,17–19^. In fact, our data suggest BQ affects mitochondrial metabolism as demonstrated by modulation of TCA cycle intermediates **(Supplementary Fig. 1d)**. Finally, in agreement with the role of DHODH in redox balance,^20^ BQ decreased the reduced:oxidized glutathione ratio (GSH/GSSG) at higher doses **(Fig. 1c)**. Overall, our data confirms sustained inhibition of DHODH-mediated metabolism within the range of selected BQ doses and timepoints.

We then used multiplexed isobaric tag-based quantitative mass spectrometry-based proteomics^14,15^ to determine the global temporal proteomic response to BQ **(Supplementary Table 2).** Principal component analysis (PCA) showed consistent sample clustering and a time-dependent shift in PC1 and PC2 (**Supplementary Fig. 2a-b**). Overall, we identified and quantified ∼8,000 proteins in PaTu-8988T and PaTu-8902 cells with a higher magnitude of changes in protein expression at 7 days compared to 24 h (**Fig. 1d-e**, **Supplementary Fig. 2c-d**). Among the consistently downregulated proteins, multiple ribonucleoproteins were decreased both acutely (**Fig. 1d, Supplementary Fig. 2c**) and long-term (**Fig. 1e, Supplementary Fig. 2d**), consistent with a role for DHODH in ribosome biogenesis^21^. In agreement with recent studies, we also identified an induction in HLA-I and proteins involved in antigen presentation (**Supplementary Fig. 2d**). DHODH inhibition upregulated expression of nucleoside importers like SLC29A1 in PaTu-8988T cells (**Fig. 1d-e**) confirming our metabolomics studies. Finally, we also identified a compensatory upregulation in GPX1 and GPX4, likely related to the increase in oxidative stress suggested by metabolomics (**Fig. 1e**).

We then performed gene set enrichment analysis (GSEA) as a strategy to identify pathways activated in response to long-term treatment with BQ (**Fig. 1f, Supplementary Fig. 2e, Supplementary Table 3)**). Functional pathway analysis identified downregulated enrichment map nodes related to glycolysis, mTOR pathway, inflammation, and TLR-related innate immunity, among others. Interestingly, upregulated nodes correlated with pathways associated with metabolic adaptations like mitochondrial translation, electron transport chain, TCA cycle, and fatty acid oxidation/sphingolipid metabolism, suggesting a compensatory response involving multiple mitochondrial-related processes (**Fig. 1f**). Next, to identify drugs with similar or inverse patterns of response to DHODHi that may be useful to overcome DHODHi-associated adaptations, we used the Connectivity Map database^22^, a resource we used in the past to define combinatorial strategies with KRAS^G12C^ inhibitors^14^. Here, we identified similar patterns of response between BQ and bromodomain, HDAC, IMPDH, or topoisomerase inhibition or HIF activation (**Supplementary Fig. 2f**).

### Defining synthetic lethalities with DHODHi by BQ-anchored CRISPR/Cas9 genetic screens

To identify synthetic lethal interactions not predicted by metabolomics or proteomics, we performed a BQ-anchored CRISPR/Cas9 genome-wide^23^ screen in PaTu-8988T cells (**Fig. 1g, Supplementary Table 4**). Next-generation sequencing followed by MAGeCK/STARS analysis highlighted genes with depleted sgRNAs in the BQ treated conditions suggesting combinatorial targets for enhanced BQ sensitivity and genes with enriched sgRNAs that may promote further resistance to BQ. Gene Ontology (GO) analysis of the top 100 genes with depleted sgRNAs overlapped with pathways identified by metabolomics and proteomics, including nucleoside salvage and regulation of steroid metabolism (**Supplementary Fig. 3a**). Furthermore, GO also identified terms not appreciated in the proteome and metabolome analyses such as “positive regulation of apoptosis intrinsic pathways” (**Supplementary Fig. 3a**). Among the significant depleted sgRNAs in the low-dose BQ screen (**Supplementary Fig. 3b**) were those targeting DHODH; however, these sgRNAs were not depleted in the high-dose BQ screen, (**Fig. 1h**) consistent with sub-maximal inhibition of BQ in the low-dose condition. Analysis of the sgRNAs consistently depleted at low and high doses of BQ (**Fig. 1h, Supplementary Fig. 3b**) confirmed nucleotide synthesis/salvage pathways (*UCK2, TK1, SLC29A1, CMPK1)* as synthetic lethalities, consistent with a compensatory upregulation in nucleotide salvage. The other most significantly depleted hits included the anti-apoptotic genes *BCL2L1* (*BCL-X_L_*) and *MCL1*, suggesting synthetic lethalities with DHODHi. Analysis of the top enriched sgRNAs (sgRNAs conferring resistance to DHODHi) identified multiple genes encoding mitochondrial complex subunits **(Supplementary Fig. 3c-d)** such as *NDUFA6, NDUFA1, NDUFS2, NDUFV1, NDUFA11* among others, which suggests Complex I inhibition may increase resistance to BQ.

The relative cellular dependency on de novo pyrimidine biosynthesis versus nucleotide salvage can shift depending on uridine availability^24,25^ and uridine levels are different in cell culture conditions versus in vivo. To account for the physiological differences in uridine levels as well as any other differences between in vitro culture conditions and in vivo conditions, we designed a focused sgRNA library to evaluate high priority hits in a BQ-anchored mini-pool in vivo CRISPR/Cas9 screen. Our custom library contained sgRNAs targeting 369 genes **(Fig. 2a)** that we prioritized based on the following criteria: 1) proteomic hits with FDA drugs available, 2) most significant hits from the CRISPR screen, including depleted and enriched guides (for balancing the library), and 3) genes that had already been linked to DHODH biology in published in vitro studies but not in in vivo studies. We first confirmed the performance of our custom library and correlation with the whole-genome screen data by testing the mini-library *in vitro* in PaTu-8988T and PANC-1 cells treated at low and high concentrations of BQ **(Fig. 2b, Supplementary Table 5).** Consistently, genes in nucleotide synthesis/salvage (*SLC29A1, UCK2, TK1, CMPK1*) were depleted across cell lines and doses **(Fig. 2c-d, Supplementary Fig. 4a-b)**. We further confirmed anti-apoptotic *BCL2L1* as a depleted gene in all cell lines. Next, PaTu-8988T cells infected with our custom library were implanted subcutaneously in the flanks of NOG mice(NOD.Cg-*Prkdc^scid^ Il2rg^tm1Sug^*/JicTac) and treated with vehicle or BQ 50 mg/kg per day for 14 days **(Supplementary Table 5)**. The top depleted hits were in agreement with some of the in vitro dependencies, and highlighted genes in nucleotide salvage (*UCK2, SLC29A1, CMPK1*) and apoptosis (*BCL2L1)* **(Fig. 2e-f)** confirming the overlap between in vitro and in vivo metabolic dependencies^7^ for our top hits **(Supplementary Fig. 4c-d)**. However, MCL1 was no longer confirmed as a hit in the in vivo screen, suggesting a selective dependency on *BCL2L1* (BCL-X_L_) and not on other anti-apoptotic proteins in vivo. Overall, our combinatorial platform identified new vulnerabilities to target in combinatorial strategies and suggests that inhibiting nucleotide salvage or anti-apoptotic proteins may synergize with DHODH inhibitors.

**Figure 2.**
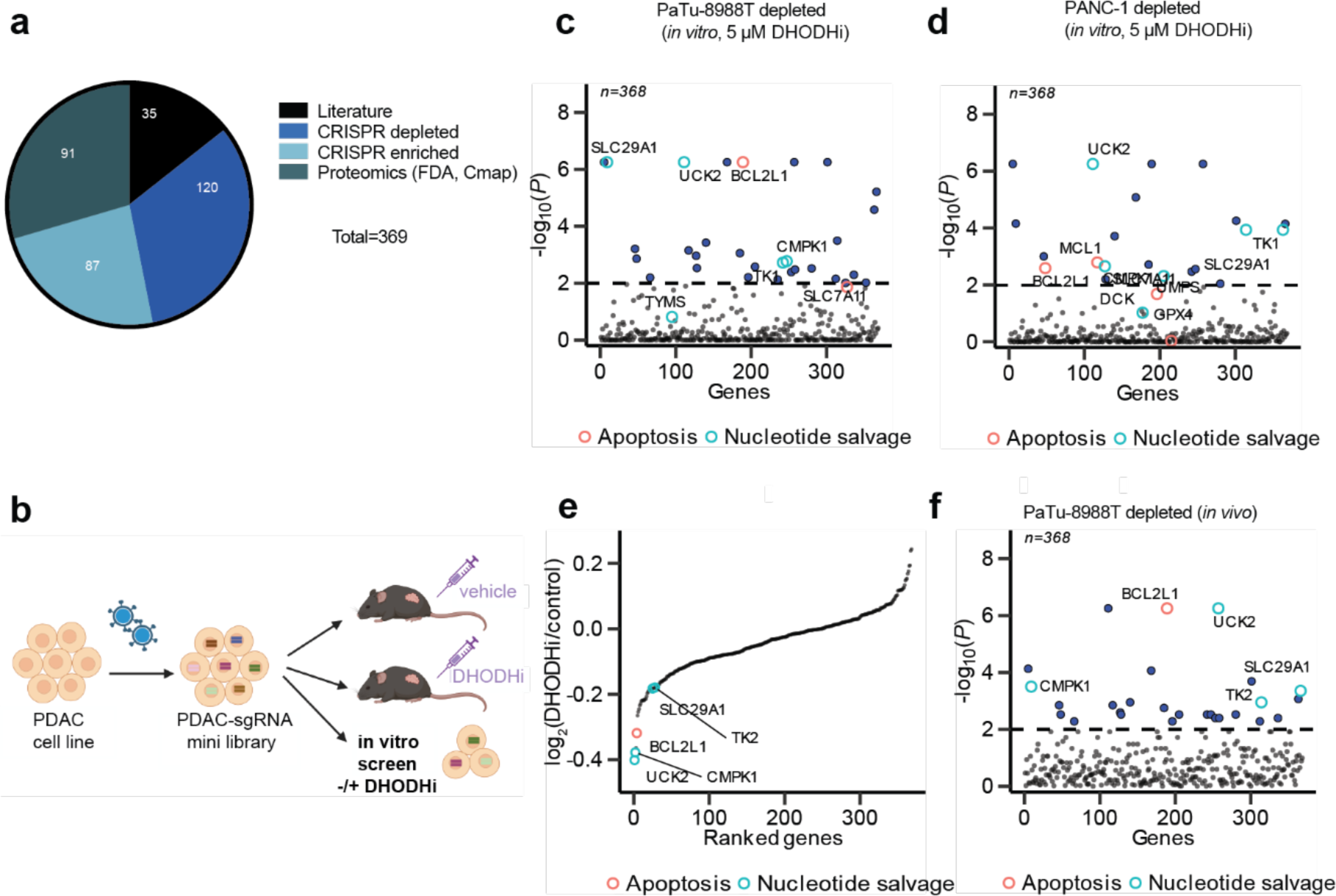
Targeted in vivo CRISPR/Cas9 loss-of-function screen identifies synthetic lethalities with BQ. **(a)** A customized CRISPR/Cas9 mini-library (369 genes, 4 sgRNAs per gene, 528 control sgRNA) was generated prioritizing proteomic hits with FDA-approved available drugs, CRISPR depleted/enriched hits, genes related to DHODH in published in vitro studies. **(b)** Schematic for in vivo and in vitro CRISPR screens with customized mini-library. PDAC cells were infected with the mini-library and implanted in a xenograft NOG mouse model (PaTu-8988T, n=10 tumors/arm). Tumors (80-100 mm^3^) were treated with either vehicle or BQ, harvested 14 days later and analyzed by NGS. PaTu-8988T and PANC-1 cells containing the sgRNA mini-library were also evaluated in in vitro screens to confirm whole genome screen and directly contrast to in vivo results. **(c-d)** Manhattan plot for depleted hits in a mini-library in vitro CRISPR screen in PaTu-8988T **(c)** and PANC-1 **(d)** cells at 5 μM BQ. Blue dots highlight significant genes (- log_10_(P)=2) in BQ *vs*. DMSO, orange circles highlight genes related to apoptosis and cyan circles highlight genes in nucleotide salvage. **(e)** Rank ordered graph of log_2_ (BQ/control) for each gene in PaTu-8988T tumors. **(f)** Manhattan plot for depleted hits in PaTu-8988T tumors. Blue dots highlight significant genes (-log_10_(P)=2) in BQ vs. vehicle, orange circles highlight genes related to apoptosis and cyan circles highlight genes in nucleotide salvage.

### Combination targeting of DHODH and BCL-X_L_ in PDAC synergistically induces apoptosis in vitro

Our CRISPR/Cas9 screens nominated inhibition of *BCL2L1*, which encodes for BCL-X_L_, as a target capable of inducing synthetic lethality in cells treated with BQ. To assess this, we evaluated the combination of BQ and DT2216^26^, a BCL-X_L_ proteolysis targeting chimera (PROTAC) that is currently being tested in Phase I clinical trials (NCT04886622). This selective PROTAC is particularly promising as it can spare platelet toxicity associated with BCL-X_L_ inhibitors^27^. Combination treatment with BQ and DT2216 demonstrated synergy in a panel of human and murine PDAC cell lines (**Fig. 3a-b, Supplementary Fig. 5a-e),** which was replicated using a distinct DHODH inhibitor, BAY-2402234 (**Supplementary Fig. 5f-g)**. We then measured cell death by PI/Annexin staining after 3 days of treatment **(Fig. 3c-d)** and real-time tracking of Annexin intensity accumulation **(Fig. 3e-f, Supplementary Fig. 5h-i).** Combination BQ and DT2216 increased apoptosis in human and murine PDAC cell lines compared to control and monotherapy and the effects were rescued by high-dose uridine and inhibition of apoptosis with Z-VAD-FMK **(Fig. 3e-f).** To assess the potential of this combination in recently described physiological media preparations that contain higher uridine levels that may decrease DHODH inhibitor sensitivity, we measured relative growth of PaTu-8902 cells in Plasmax media that contains 3 µM uridine that more closely matches levels in human plasma in comparison to DMEM media preparations **(Supplementary Fig. 5j-m)**. Consistent with previous studies, the anti-proliferative effects of BQ and BAY-2402234 were less striking in closer-to-physiological conditions. Nevertheless, BQ and DT2216 significantly inhibited growth compared to single treatments **(Supplementary Fig. 5j-m)**, suggesting that targeting de novo pyrimidine biosynthesis under physiologic uridine microenvironmental conditions still potentiates to BCL-X_L_ targeting and may maintain combination efficacy in vivo.

**Figure 3.**
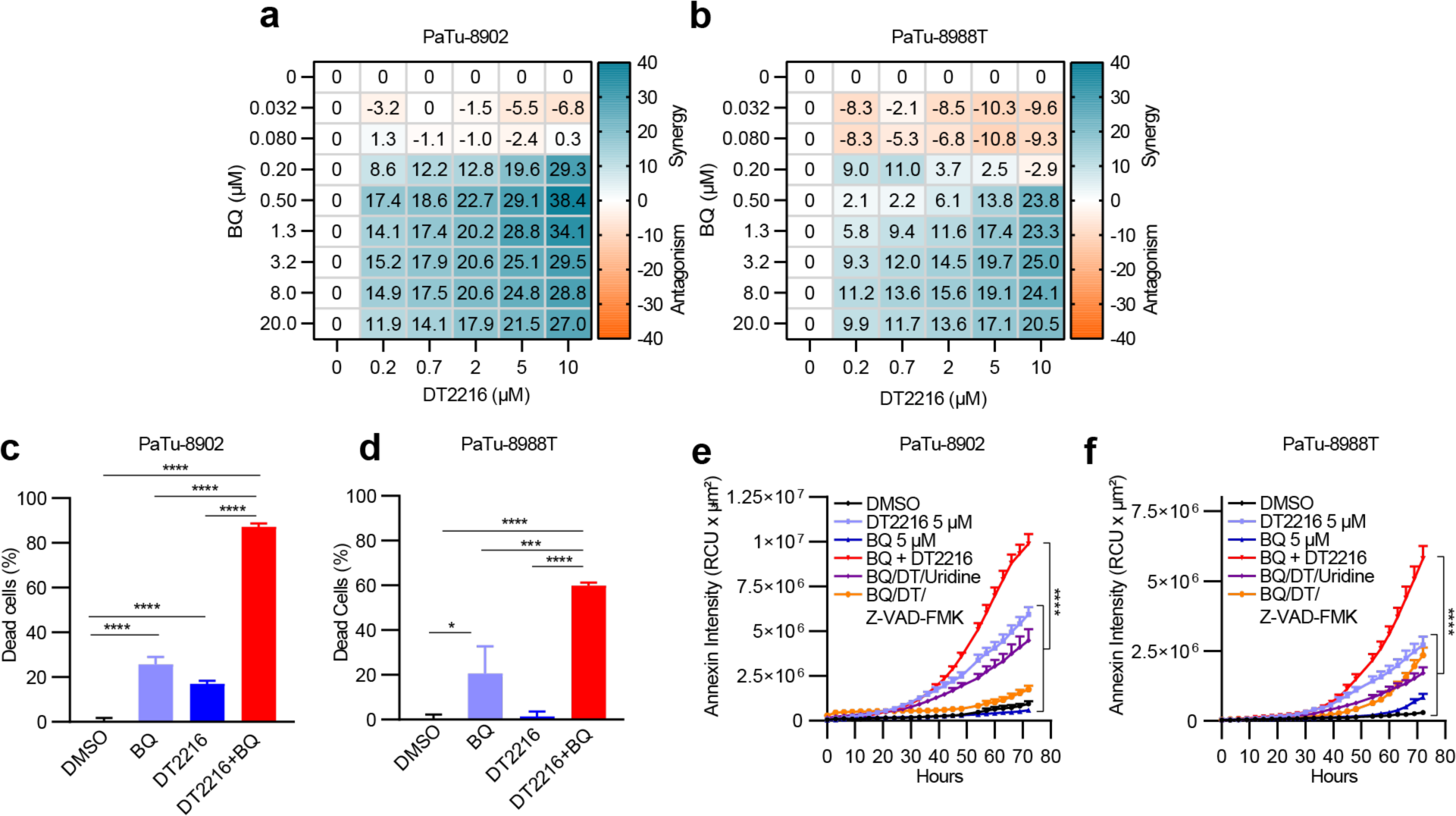
Combination of BQ and DT2216 synergistically induces apoptosis in pancreatic cancer cell lines. **(a-b)** Synergy score heatmaps of combination treatment with BQ and DT2216 in PaTu-8902 and PaTu-8988T cells. Synergy score between the two drugs was calculated using the HSA model implemented in SynergyFinder (antagonism: <− 10; additive effect: from − 10 to 10; synergistic effect: >10). Experiments performed in biological triplicate, data shown as mean HSA score of three technical replicates of one representative experiment. **(c-d)** Cell death analyzed by flow cytometry after treatment with DMSO, DT2216 (2 µM) or BQ (5 µM) alone or in combination for 72 hours. Error bars represent s.d. of three technical replicates (representative of three experiments). **(e-f)** Real-time Annexin V fluorescence signal accumulation in PaTu-8902 and PaTu-8988T. Cells were labeled with Annexin V Red Dye and treated with the indicated concentrations of BQ or DT2216 alone or in combination or BQ + DT2216 with uridine (100 µM) or Z-VAD-FMK (50 µM) for 72 hours. (RCU: Red Calibrated Unit). Error bars represent s.d. of three technical replicates (representative of two experiments). For all panels, significance determined with ordinary one-way ANOVA test. **p < 0.01, ***p < 0.001, ****p < 0.001.

### BQ modulates the balance of BCL-2 family proteins to induce a pro-apoptotic shift

Previous studies have shown that BCL-X_L_ expression increased from the preinvasive pancreatic intraepithelial (PanIN) stage to invasive PDAC in mouse models and patient tumors^28^. Importantly, compared to other anti-apoptotic proteins, higher levels of BCL-X_L_ were observed in 90% of tumors from PDAC patients^29^. Monotherapy targeting of BCL-X_L_ has modest effects in solid tumors; however, combinations of BCL-X_L_ with gemcitabine and nab-paclitaxel or MEK inhibition have been reported to enhance PDAC anti-tumor efficacy^30-31^. Our data suggests DHODH inhibition sensitizes PDAC to apoptosis inducers targeting BCL-X_L_. As the apoptotic cascade is a complex process regulated by multiple BCL-2 family proteins, we investigated how DHODH may rewire the apoptosis pathway in PDAC. To assess if BQ modulates the overall apoptotic priming as well as dependencies on specific pro-survival proteins in PDAC cells, we performed flow cytometry-based BH3 profiling^32^ **(Fig. 4a-b)**. At basal levels, PaTu-8988T was more primed than PaTu-8902 by response to BIM and PUMA peptide, as indicated by increased cytochrome c release in response to pro-apoptotic BIM, BID, and PUMA BH3 peptides, indicating that PaTu-8988T will more readily undergo apoptosis in response to damage or stress than PaTu-8902^33,34^. Although BQ did not significantly alter the priming of either cell line, this treatment increased dependency on BCL-X_L_ and BCL2 in both cell lines as indicated by the increased response to BAD and HRK peptides; however, BQ did not significantly alter their priming status. Next, to understand how BQ changes the dependency to different pro-apoptotic proteins, we quantified expression of the BCL-2 family proteins after BQ treatment. Proteomic quantification of proteins in the apoptotic pathways demonstrated BQ decreased expression of anti-apoptotic proteins (BCL-X_L_, MCL1) while increasing expression of pro-apoptotic proteins BAX, BAK, BID, NOXA and PUMA (**Supplementary Fig. 6a-b**). Western blot analyses confirmed the pro-apoptotic shift induced by BQ with decreases in anti-apoptotic proteins (BCL-X_L_, MCL1, BCL2) and increased levels of cleaved-PARP and pro-apoptotic proteins BIM and PUMA (**Fig. 4c**). Next, we evaluated the BCL-2 protein profile induced by DT2216 alone (**Fig. 4d, Supplementary Fig. 6c**) or in combination with BQ (**Fig. 4e, Supplementary Fig. 6d**). DT2216 efficiently degraded BCL-X_L_ in the range of doses tested (0.1-5 μM) in human and murine PDAC cells. Of note, BCL-X_L_ degradation induced compensatory increases in MCL-1 and BCL-2 (**Fig. 4d, Supplementary Fig. 6c**). Finally, combination of BQ and DT2216 significantly increased PARP cleavage and BIM compared to single treatment (**Fig. 4e, Supplementary Fig. 6d**). Together, our data suggests BQ enhances the effects of BCL-X_L_ degraders by shifting the balance between anti-apoptotic and pro-apoptotic proteins to induce cytotoxicity.

**Figure 4.**
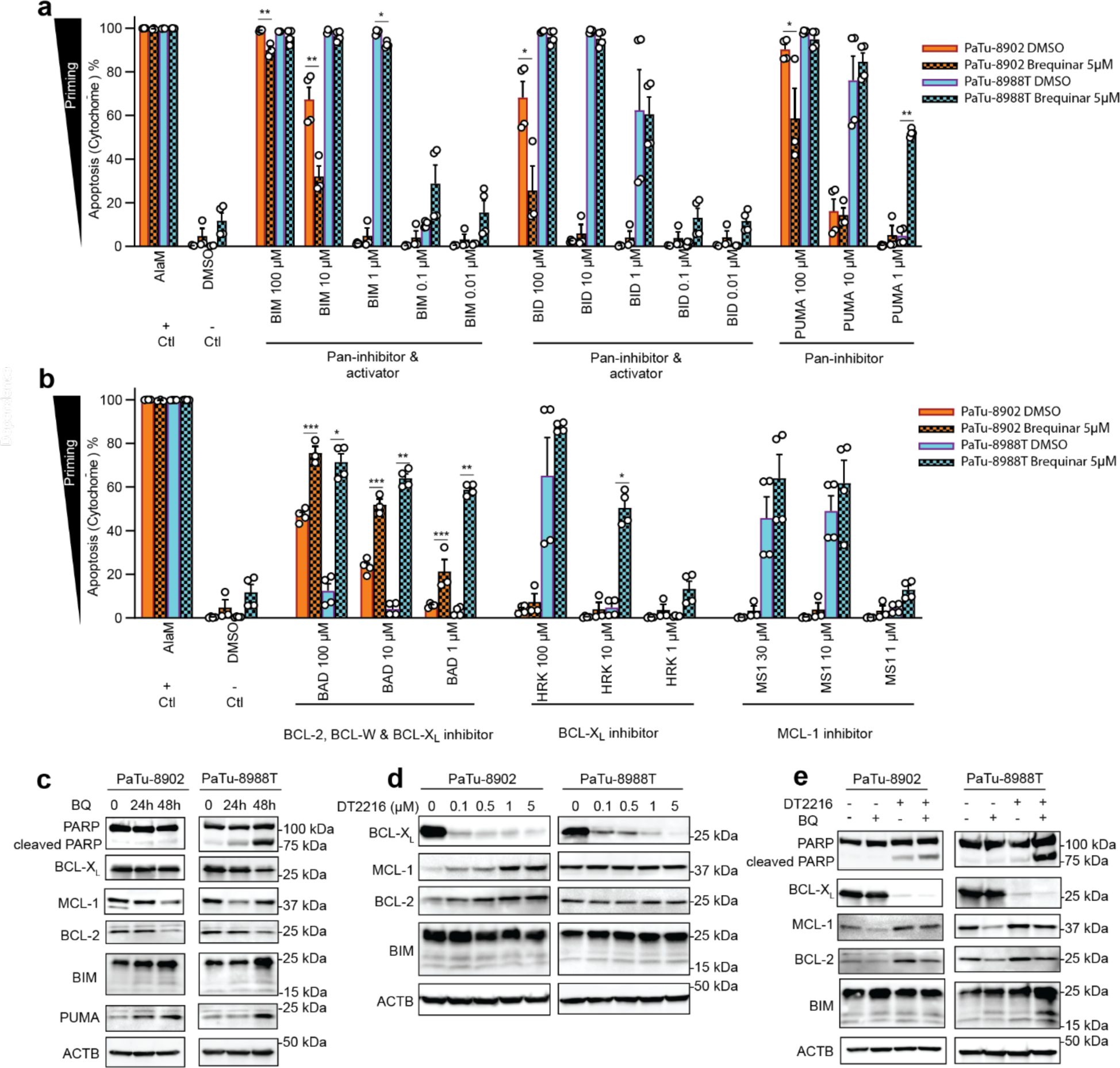
BQ modulates levels of BCL-2 family proteins and enhances sensitivity to BCL-X_L_ PROTAC-based degradation. **(a-b)** Flow cytometry-based BH3 profiling of PaTu-8902 and PaTu-8988T cells treated with BQ for 7 days shows priming **(a)** and the specificity of BCL-2 protein family apoptotic dependency **(b)**. Apoptosis is measured by Cytochrome c release (y-axis, see meethods). Alamethicin (AlaM) was used as positive control for apoptosis induction. DMSO was used as a negative control. BIM, BID and PUMA BH3 peptides can inhibit all the anti-apoptotic proteins and BIM and BID can also directly activate BAX and BAK. These conditions are used to determine the overall level of priming in a cell. BAD BH3 peptide can inhibit BCL2, BCL-W and BCL-X_L_ and indicates dependence on these proteins while HRK and MS1 peptides selectively inhibit BCL-X_L_ and MCL1, respectively. Error bars represent SEM of two independent experiments, significance determined with t-test. *p < 0.05, **p < 0.01, ***p < 0.001. **(c)** Immunoblot analysis of BCL-2 family proteins as indicated in lysates from PaTu-8902 and PaTu-8988T cells treated with 5 µM BQ for 24 and 48 hours. **(d)** Immunoblot analysis of BCL-X_L_, BCL2 and MCL1 levels in lysates from PaTu-8902 and PaTu-8988T cells treated with the indicated concentrations of DT2216 for 16 hours. (**e)** Immunoblot analysis of BCL-2 family proteins in lysates from PaTu-8902 and PaTu-8988T cells treated with 5 µM BQ or 5 µM DT2216 alone or BQ in combination with DT2216 for 24 hours.

### BQ promotes synergistic responses selectively with BCL-X_L_ inhibitors

Given the broad effects of BQ on multiple BCL-2 family proteins and identification of MCL1 sgRNAs as depleted in the in vitro CRISPR/Cas9 screen, we assessed if combination with other BCL-2 protein family inhibitors similarly synergized with BQ. Cell viability assays confirmed synergistic effects of BQ and BCL-X_L_ inhibition (DT2216 or A-1331852) using either DT2216 (**Fig. 5a, d**) or BQ as an anchor (**Supplementary Fig. 7a-d**). Interestingly, inhibitors against other BCL-2 family proteins such as AZD-5591, an MCL-1 inhibitor (**Fig. 5b, e**) or venetoclax, a BCL-2 inhibitor (**Fig. 5c, f**), demonstrated no synergy with BQ and in fact demonstrated protective effects at high concentrations. Overall, our data suggest the BQ-induced pro-apoptotic shift only induces sensitivity to BCL-X_L_ inhibition and no other BCL-2 family members. In fact, BCL-X_L_ showed strong PDAC dependency compared with MCL1 and BCL2 across 45 human lines in the Cancer Dependency Map (DepMap), a library of genome-wide CRISPR loss-of-function screens^35,36^ (**Supplementary Fig. 7e**). Querying DepMap highlighted PaTu-8902 as the cell line with highest level of BCL-X_L_ expression and dependency on BCL-X_L_ across the 6 human PDAC lines we tested (**Supplementary Fig. 7f**), which correlated with the basal BCL-X_L_ level detected by immune blot (**Supplementary Fig. 7g**) and the strong response to BQ and DT2216 (**Fig. 3a,c**). Our results demonstrate that BQ promotes synergistic responses exclusively with BCL-X_L_ inhibitors and not inhibitors of BCL2 or MCL1, which may be because BQ increases cell dependency on BCL-X_L_, a critical mediator of PDAC cell survival^37^.

**Figure 5.**
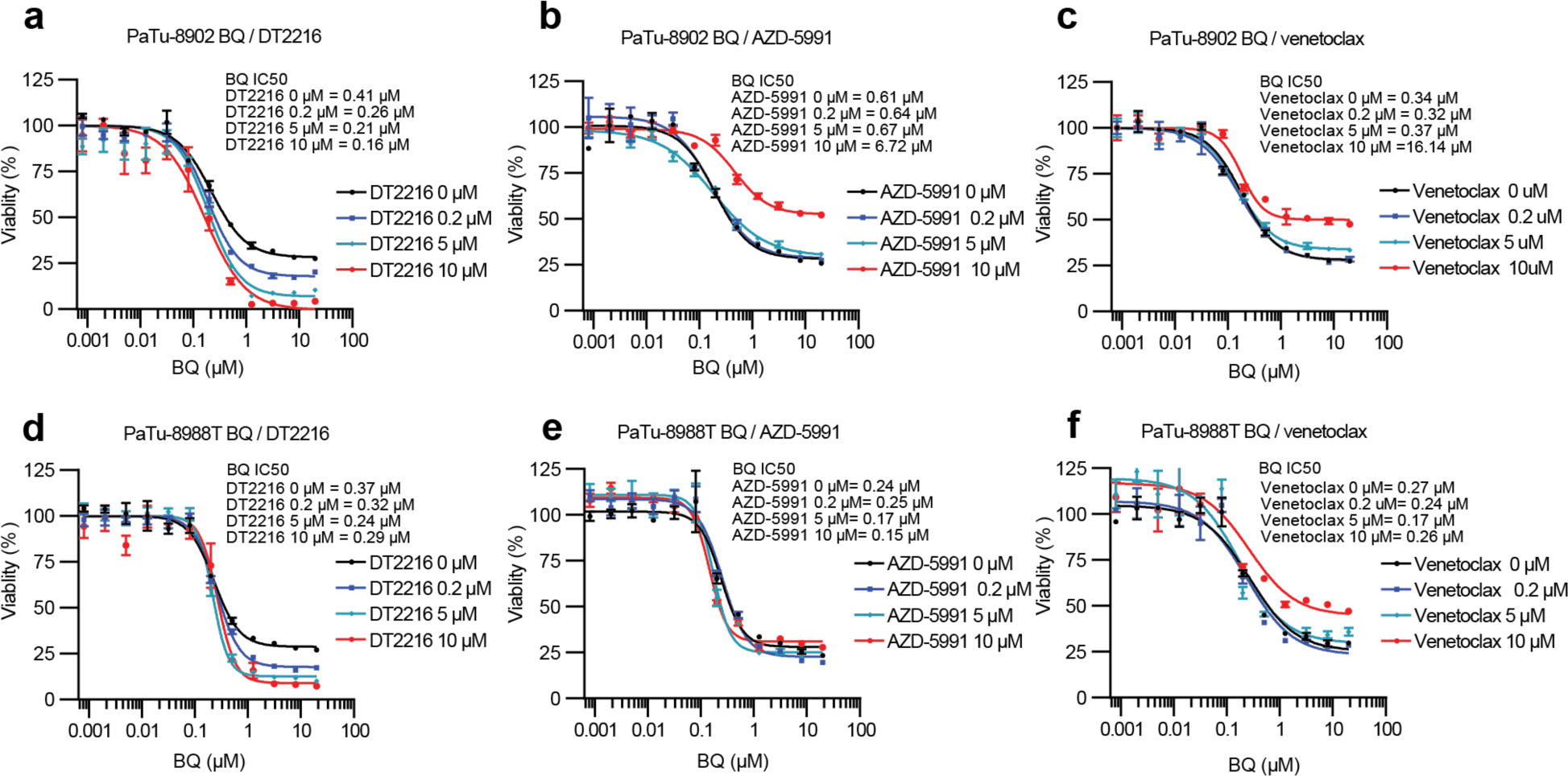
Synergistic effects with BQ are specific to BCL-X_L_ inhibition. **(a-f)** Percentage relative growth of PaTu-8902 and PaTu-8988T cells after treatment with increasing concentrations of BQ with DT2216 (BCL-X_L_ PROTAC), AZD-5991 (MCL1 inhibition) and Venetoclax (BCL2 inhibition) respectively for 5 days. IC50 values are shown for a representative experiment out of three independent experiments.

### Combination targeting of DHODH and BCL-X_L_ inhibit PDAC growth in patient-derived organoids and in vivo models of PDAC

To bridge the gap between our results in established PDAC monolayer culture systems and the potential in vivo effects of this combination, we tested the combination of BQ and DT2216 in a collection of patient-derived PDAC organoids that can more closely reflect in vivo tumor phenotypes and response to therapy (**Supplementary Table 6**). Drug sensitivity and synergy in patient-derived organoids was consistent with our results in established PDAC cell culture models (**Fig. 6a-d**), suggesting that combination targeting of DHODH and BCL-X_L_ may inhibit tumor growth in tumor models of PDAC. Interestingly, three of the organoids used were KRAS^G12D^ mutant but a priori resistant to pharmacologic KRAS^G12D^ inhibition suggesting BQ and DT2216 may be useful even for KRAS inhibitor resistant tumors. Indeed, a KRAS^G12D^-mutant PDAC cell line exposed to escalating doses of MRTX1133 to the point of resistance maintained sensitivity to the combination of DHODH and BCL-X_L_ inhibition (**Supplementary Fig. 8a**).

**Figure 6.**
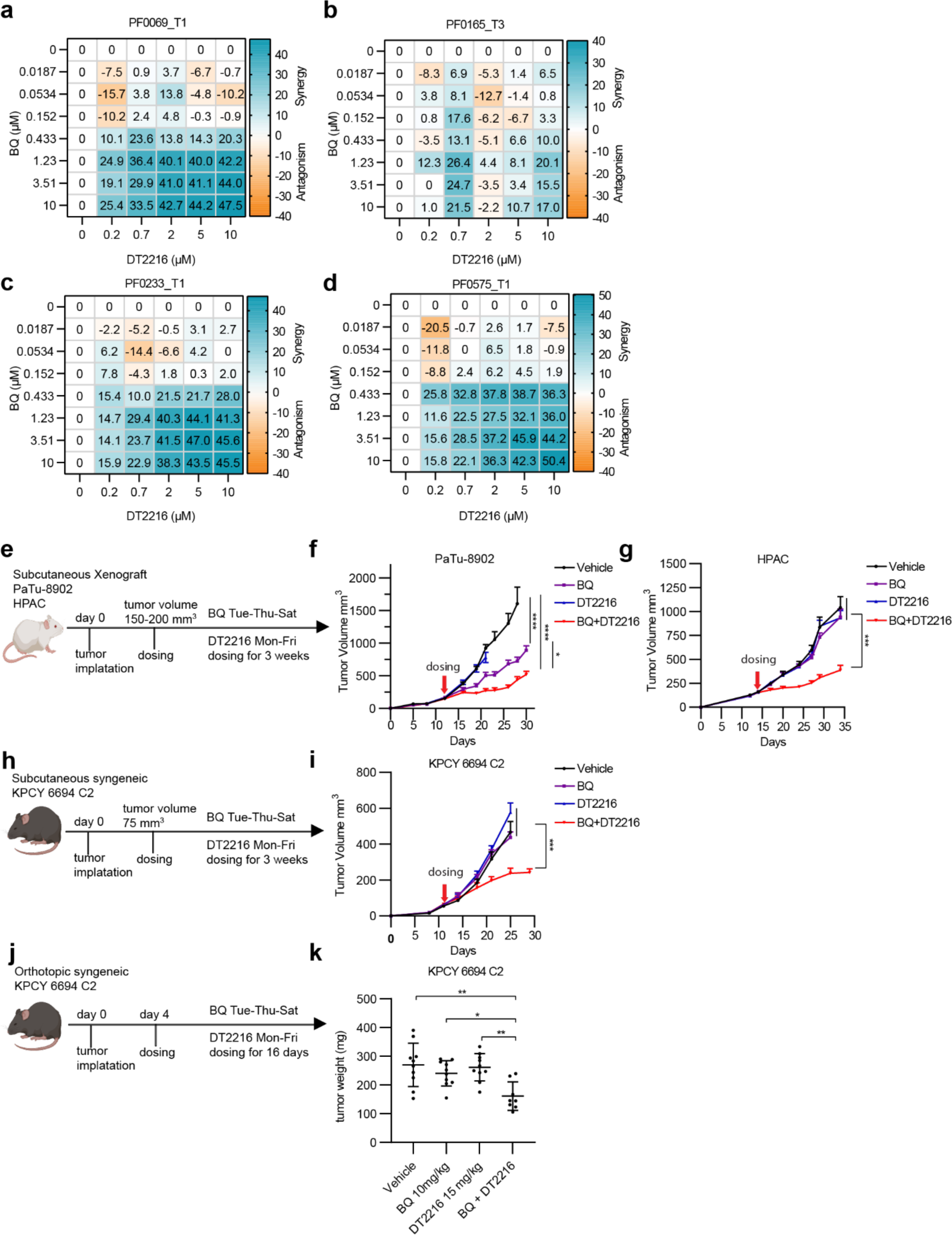
DT2216 increases the antitumor efficacy of BQ in patient-derived organoids and PDAC mouse models. **(a-d)** Synergy score heatmaps of patient-derived organoids treated with DT2216 and BQ. The synergy score was calculated using the HSA model implemented in SynergyFinder. The experiments were repeated two independent times, and data are shown as mean HSA score of three technical replicates from one experiment. **(e)** Experimental design of PaTu-8902 and HPAC xenograft flank tumor studies in immune deficient mice. **(f-g)** Tumor growth curves from PaTu-8902 (**f**, n=13 per arm) and HPAC (**g**, n=7 per arm) xenograft mouse models treated with vehicle, BQ, DT2216, or BQ in combination with DT2216 for 3 weeks. **(h)** Experimental design of syngeneic flank allograft study. **(i)** Tumor growth curve of KPCY 6694 C2 syngeneic mouse model treated with vehicle (n=12), BQ (n=13), DT2216 (n=10) or BQ in combination with DT2216 (n=10) for 3 weeks. **(j)** Experimental design of syngeneic orthotopic tumor study. **(k)** Tumor weight of KPCY 6694 C2 tumors (n=10 per arm) after treatment with vehicle, BQ, DT2216, or BQ in combination with DT2216 for 16 days. For flank-based tumor in vivo studies, tumors were measured twice a week. Error bars represent ± SEM. Statistical significance was determined by ordinary one-way ANOVA test. *p < 0.05, **p < 0.01, ***p < 0.001.

To further assess this combination as a potential therapeutic strategy for PDAC, we first evaluated for maximum tolerated doses of the combination of BQ and DT2216 in non-tumor bearing animal models. We noted that full dose BQ in combination with published regimens of DT2216 was associated with dose-limiting hematologic toxicity, in line with both DHODH inhibition and BCL-X_L_ inhibition having effects on the bone marrow. We established BQ at 10 mg/kg three times per week and DT2216 at 15 mg/kg two times per week as a tolerable regimen (**Supplementary Fig. 8b**) with consistent BCL-X_L_ degradation in the pancreas and liver (**Supplementary Fig. 8c**) by DT2216 alone or BQ and DT2216 in combination. We next investigated the efficacy of BQ and DT2216 using PaTu-8902 and HPAC cell line-derived xenograft tumor models. DT2216 alone had minimal effects on tumor growth in vivo whereas BQ had some monotherapy efficacy in PaTu-8902 but none in HPAC xenografts. However, the combination of BQ and DT2216 led to a significant inhibition of tumor growth (**Fig. 6e-g**), even in HPAC xenografts where there was no single agent efficacy. Next, we examined the anti-tumor activity of the combination in an allograft KPCY C57BL/6 mouse that allows for evaluation in an immune competent setting. The poorly immunogenic murine PDAC cell line KPCY 6694 C2^38^ was implanted subcutaneously **(Fig. 6h-i)** and allowed to establish tumors (75 mm^3^) prior to dosing. Similar to HPAC there was minimal single agent efficacy but a significant growth delay with the combination. To model the pancreatic microenvironment where nutrient supply may be differential compared to the subcutaneous environment, we orthotopically implanted KPCY 6694 C2 cells^39^ and allowed tumors to establish prior to dosing **(Fig. 6j-k)**. The BQ and DT2216 combination significantly suppressed the growth of orthotopic KPCY 6694 C2 tumors (**Fig. 6k**). The combination treatment was well tolerated in the subcutaneous tumor mouse models (**Supplementary Fig. 8d-f**), with no significant change in mouse body weight after 3 weeks whereas, we observed a modest weight drop in the orthotopic tumor model (**Supplementary Fig. 8g**). While hematologic parameters were largely unaffected in NSG mice, the BQ and DT2216 combination demonstrated modest anemia and thrombocytopenia in C57BL/6 mice (**Supplementary Table 7-8**). To confirm the on-target effects of DT2216, we evaluated BCL-X_L_ degradation in tumors by immunoblotting. DT2216 decreased tumor BCL-X_L_ levels in xenograft models and to a lesser degree in syngeneic models (**Supplementary Fig. 8h-o)**, which may, in part, explain reduced tumor growth inhibition in the syngeneic experiments. Finally, we found higher levels of cleaved caspase 3 in PaTu-8902 flank tumors, indicating the drug combination induces apoptosis in vivo (**Supplementary Fig. 8p)**. Overall, we propose combinatorial targeting of DHODH and BCL-X_L_ as a therapeutic strategy with potential clinical translatability to restrict PDAC growth.

## DISCUSSION

There is renewed interest in repurposing DHODHi as anti-cancer agents given the importance of de novo pyrimidine biosynthesis as a cancer dependency. While DHODHi failed as a monotherapy in previous phase I clinical trials for solid tumors^9,12^, an important caveat of these studies was the use of intermittent dosing patterns, similar to cytotoxic chemotherapy regimens, that allow for recovery of pyrimidine pools and may be insufficient to consistently deplete pyrimidines in the tumor^4^. New trials are currently evaluating the efficacy of more potent DHODHi in hematological malignancies (NCT04609826 and NCT02509052) given their ability to induce differentiation of diverse AML subtypes^40,41^. Targeting DHODH in solid tumors is also an exciting avenue; however, this will likely rely on identification of combinatorial strategies to improve monotherapy responses.

DHODH inhibition has been previously linked with apoptosis induction in different types of cancer including hematological malignancies^42,43^, colorectal cancer^44^, neuroblastoma^45^, renal cell carcinoma^46^ and glioma^47,48^. For example, silencing of DHODH or DHODH inhibition by high-dose BQ sensitized the small cell lung cancer U1690 cell line to TRAIL-induced apoptosis^49^. Replication stress induced by DHODH inhibition has also been shown to contribute to its role in apoptosis induction^40^. However, in pancreatic cancer it is unclear how DHODH inhibition can regulate apoptosis sensitivity. It has been reported that teriflunomide, a DHODH inhibitor, induces apoptosis in pancreatic cancer by directly targeting the apoptotic regulator PIM-3 serine-threonine kinase^50^. Our results suggest that BQ-induced pyrimidine depletion sensitized PDAC cells to apoptosis by decreasing anti-apoptotic proteins and increasing pro-apoptotic proteins. A similar profile of anti-apoptotic protein alteration in response to gemcitabine was previously demonstrated suggesting a conserved response to nucleotide metabolism targeting^32^; however, the specific mechanism by which DHODH inhibition induces these alterations is unclear and warrants future study. Functionally, the induced alteration in anti-apoptotic proteins whether by DHODH inhibition or gemcitabine conferred increased sensitivity specifically to BCL-X_L_ targeting.

The specificity of enhanced sensitivity to BCL-X_L_ targeting conferred by DHODH inhibition in PDAC appears be ingrained in PDAC tumor physiology at baseline. Previous studies have shown increased expression of BCL-X_L_ with the progression of pancreatic neoplasia from pancreatic intraepithelial neoplasia (PanIN)-1 to PDAC, while MCL1 slightly increased and BCL2 did not change^28^. Furthermore, BCL-X_L_ but not MCL1 or BCL2 scores as a PDAC cell line dependency in DepMap data (**Supplementary Fig. 7e**). The importance of BCL-X_L_ in PDAC physiology was further highlighted by Stanger and colleagues who demonstrated that BCL-X_L_ enforces a slow-cycling state allowing survival of quiescent PDAC cells present in the austere nutrient and oxygen deprived PDAC tumor microenvironment^31^. These findings suggest an extra-reliance on BCL-X_L_ in PDAC in comparison to MCL1 and BCL2 and may in part explain the selective synergy of DHODH inhibition with BCL-X_L_ inhibition.

Our DHODH inhibition multi-omic data also demonstrate that activation of nucleoside salvage is a compensatory adaptation to DHODHi and suggests a combination therapeutic strategy. However, our studies present some limitations. First, our initial screens were performed in standard culture conditions, which differ in the amount of uridine and other metabolites available in comparison to the in vivo tumor metabolic milieu. Uridine levels in tumor interstitial fluid of subcutaneous and orthotopic PDAC tumors can be as high as 30-50 μM (10 times more than plasma concentrations). In fact, activation of nucleoside salvage is likely a contributor to the poor anti-tumor effects of monotherapy DHODHi in the uridine rich TME^51^. Indeed, exogenous uridine rescues growth defects in DHODHi-treated cells cultured in standard media conditions^24,25,52^ and partially rescues our combinatorial treatment **(Fig. 3)**. To account for differences in metabolite availability in standard DMEM versus in vivo^51^, cells can be cultured in media that mimics the physiological metabolic conditions of adult human plasma (e.g. Plasmax, HPLM) or the PDAC TME. Prior studies have shown that cancer cells grown in HPLM not only utilize uridine for nucleoside salvage, they also suppress de novo pyrimidine synthesis by uric acid-mediated inhibition of UMPS activity, decreasing sensitivity to pyrimidine synthesis inhibitors like 5-FU^25^. To account for these known differences in standard DMEM media preparations, we performed a secondary CRISPR/Cas9 screen in vivo in the context of a fully formed tumor. Interestingly, both our in vitro and in vivo CRISPR screens highlighted the nucleoside salvage pathway as a co-dependency with DHODH inhibition. Despite likely differences in the relative usage of de novo pyrimidine biosynthesis versus nucleotide salvage pathways in vitro versus in vivo at baseline, DHODH inhibition in vivo still demonstrated synergy with nucleotide salvage. This suggests that despite the high levels of uridine in the TME there remains a metabolic consequence of DHODH inhibition in vivo in PDAC tumors, namely pushing tumors to rely more heavily on nucleoside salvage. One therapeutic strategy suggested by dual reliance on pyrimidine synthesis and salvage pathways in PDAC is combination targeting of pyrimidine synthesis and salvage. Indeed, combination BQ and gemcitabine has been described as a potential combinatorial strategy in PDAC with tolerable in vivo toxicity^53^. Our CRISPR screen identified SLC29A1, a nucleoside importer, as an in vivo co-dependency, in agreement with previous studies showing synergistic in vitro efficacy^16^. However, to date there are no potent, selective, and in vivo capable inhibitors available for SLC29A1. Future studies are required to assess the feasibility of this combination since systemic dual targeting of nucleoside import and de novo pyrimidine synthesis may result in systemic toxicities affecting cells with high biosynthetic needs in the bone marrow and intestine as well as most non-tumor cells that mostly rely on salvage.

Combinatorial targeting of DHODHi with BCL-X_L_ was effective through multiple cell culture conditions (DMEM, Plasmax), in vivo models of PDAC and even KRAS inhibitor-resistant organoid systems, suggesting a conserved combinatorial effect in different metabolic environments and a potential use of this strategy to sensitize KRASi-resistant PDAC cells. BCL-X_L_ is a well-demonstrated cancer target; however, dose-limiting thrombocytopenia limits the use of BCL-X_L_ inhibitors clinically^54^. DT2216 is a BCL-X_L_ PROTAC that targets BCL-X_L_ to the Von Hippel-Lindau (VHL) E3 ligase for degradation^26^. Given the lower expression of VHL in platelets, DT2216 is less able to degrade BCL-X_L_ thereby reducing thrombocytopenia^26^. No toxicity was reported when DT2216 was combined with gemcitabine in PDAC^32^ or AZD8055 in small-cell lung cancer^55^. Similarly, we did not observe weight loss or thrombocytopenia in our NSG xenograft model (**Supplementary Fig. 8d and Supplementary Table 8**). However, the combination of BQ and DT2216 caused modest anemia and thrombocytopenia in the C57BL/6J model (**Supplementary Table 7**). One strategy to mitigate potential hematologic side effects of the BQ and DT2216 combination is administration of recombinant G-CSF (filgrastim, pegfilgrastim) and romiplostim, common supportive treatments administered with FOLFIRINOX chemotherapy^51^. Although G-CSF induced neutrophils can suppress T cell activity,^56^ we have shown that the combination of BQ and DT2216 does not require adaptive immunity. Further advances in targeting BCL-X_L_ may improve tolerability as well. New PROTAC agents against BCL-X_L_ are currently being developed in order to increase BCL-X_L_ selectivity and decrease toxicity^57–60^. Targeting BCL-X_L_ to tumors via an antibody–drug conjugate (ADC) approach is also being evaluated to minimize non-tumor tissue effects^61^. ABBV-637, an antibody-drug conjugate (ADC) consisting of a monoclonal antibody directed against the epidermal growth factor receptor (EGFR) conjugated to an inhibitor of BCL-X_L_, and ABBV-155, an ADC composed of a monoclonal antibody against the immunoregulatory protein B7-homologue 3 (B7-H3, CD276) conjugated to a BCL-X_L_ inhibitor are now in Phase 1 clinical trials for lung cancer treatment (NCT04721015 and NCT03595059). Overall, we have designed a multi-omics approach that allows for discovery of drug adaptation mechanisms. We nominate DHODHi and BCL-X_L_ as a potential therapeutic strategy in PDAC.

## METHODS

### Cell Culture

PaTu-8988T and PaTu-8902 cells were obtained from DSMZ, PANC-1, HPAC, Panc 02.03 and HEK-293T cells were from ATCC and PK-1 and KP-4 were from RIKEN Cell Bank. The murine KPCY 6499 C4 and KPCY6694 C2 cell lines were from Kerafast. Cell lines were maintained in a centralized cell bank and authenticated by assessment of cell morphology as well as short tandem repeat fingerprinting. Cells were routinely tested for Mycoplasma contamination using PCR and were negative for Mycoplasma. After thawing, cell lines were cultured for no longer than 30 days. Cell lines were maintained at 37°C with 5% CO2 and grown in DMEM or RPMI 1640 supplemented with 10% FBS and 1% penicillin/streptomycin, unless otherwise specified. Plasmax™ (CancerTools, 156371) was supplemented with 2.5% FBS and 1% penicillin/streptomycin before use. HPLM (Gibco, A4899101) was supplemented with 10% dialyzed FBS (Gibco, A3382001), RPMI 1640 1X Vitamins (MCE, R7256) and 1% penicillin/streptomycin before use. Organoid cultures were established and maintained according to previously described techniques^62^ In brief, tumor cells from patients after dissociation were cultured in 3-dimensional (3D) Growth-factor Reduced Matrigel (Corning), added with human complete organoid medium containing Advanced DMEM/F12 (GIBCO), 10 mM HEPES (GIBCO), 1x GlutaMAX (GIBCO), 500 nM A83-01 (Tocris), 50 ng/mL mEGF (Peprotech), 100 ng/mL mNoggin (Peprotech), 100 ng/mL hFGF10 (Peprotech), 10 nM hGastrin I (Sigma), 1.25 mM N-acetylcysteine (Sigma), 10 mM Nicotinamide (Sigma), 1x B27 supplement (GIBCO), RSPONDIN-1 conditioned media 10% final, WNT3A conditioned media 50% final, 100 U/mL penicillin/streptomycin (GIBCO), and 1x Primocin (Invivogen), and maintained at 37°C in 5% CO2. For proliferation assays, organoids were dissociated with TrypLE Express (GIBCO) before re-seeding into fresh Matrigel and culture medium.

### Cell doubling

To assess the effects of different doses of Brequinar in PDAC cells, 300,000 cells were plated in 6 cm plates. On day 1, BQ was added at 0.5, 1 or 5 μM and cells were counted on days 3, 6, 9 and 12. Number of cells were quantified by flow cytometry (Beckman Coulter Cytoflex).

### Metabolomics

Steady state metabolomics experiments were performed as previously described^14^. PaTu-8988T cells were plated and medium was refreshed next day with BQ/DMSO. Media was refreshed 2h before metabolite collection at 24h or 7 days treatment. Experiments were performed in 25 mM glucose/4 mM glutamine. Metabolite fractions were normalized to cell number in a parallel 6 cm plate.

### Quantitative Proteomics

Mass spectrometry–based proteomics was performed as previously described^15^. Cells were lysed using 8 M urea, 200 mmol/L EPPS, pH 8.5 with protease inhibitors. 100 μg of protein extracts were reduced using TCEP and alkylated with 10 mmol/L iodoacetamide followed by chloroform/methanol precipitation. Protein pellets were digested overnight with Lys-C and trypsin digested the next day. Peptides were labeled using 200 μg of TMT reagent (20 μg/ μl). To equalize protein loading, a ratio check was performed by pooling 2 μg of each TMT-labeled sample. Pooled TMT-labeled peptide samples were fractionated by basic-pH reverse-phase HPLC. Samples were desalted using StageTips prior to analyses using LC-MS/MS/MS. All mass spectrometry data were acquired using an Orbitrap Lumos mass spectrometer in line with a Proxeon NanoLC-1200 UHPLC system. All acquired data were processed using Comet^63^ and a previously described informatics pipeline^64^. Spectral searches were done using fasta-formatted databases (Uniprot Human, 2020, or Uniprot Mouse, 2020). Protein quantitative values were normalized so that the sum of the signal for all proteins in each channel was equal to account for sample loading.

### Brequinar-anchored CRISPR/SpCas9 genetic screen

The all-in-one version of the Human CRISPR knockout Pooled Library (Addgene #73179) was applied for in vitro genome-wide loss-function screen in PaTu-8988t cells^23^. The distribution degree of all guides in amplified library were assessed via NGS to ensure the Gini index is less than 0.1. 64 μg of pooled plasmid was produced from 100 million 293T cells using Lipofectamine 3000 according to manufacturer’s standard protocol. The molar ratio of pooled plasmid, psPAX2 (Addgene #12260) and pMD2.G (Addgene #12259) was 2.5:2:1. Culture media was refreshed 24 h after transfection, and virus-containing media were collected at 48 h and 72 h. Next, virus-containing media were pooled and filtered through a 0.45 μm filter. To determine viral efficiency, dilutions of virus were mixed with 1 million PaTu-8988T cells and cells were incubated in complete culture media containing 4 μg/ml polybrene and spun at 1,850 rpm at 25 °C for 2 hours. Cells were incubated at 37 °C for 24 h followed by selection with 1 μg/ml puromycin. After a 4-day puromycin selection, cell numbers were counted, and virus efficiency was calculated as puromycin-selected well / no-puromycin well. In the full-scale experiment, 108 mL virus was used to infect 180 million cells via the plate spin infection method, and then incubated with 1 μg/ml puromycin for 5 days. The final infection ratio was 23%. Cells were amplified in complete media until 400 million cells were obtained. Cells were then divided into three arms (40 million cells as biological triplicates in each arm): DMSO, 0.5 μM BQ (low dose) and 5 μM BQ (high dose). Cells were split every 3 days and treated with BQ for 14 days in total. Cells were collected and genome DNA was extracted via Qiagen Blood & Cell Culture DNA Maxi Kit (#13362). Guide RNA sequences were amplified using Illumina sequencing specific primer sets with NEB Q5 High-Fidelity DNA polymerase (#M0491L). Amplified amplicon sequencing was performed using next-generation sequencing (NGS) at 500x depth of each guide RNA by Novogene. Acquired NGS reads were analyzed via the MAGeCK tool^65^. Candidate genes were selected based on MAGeCK’s output, considering both effect sizes and statistical significance.

### In vitro and in vivo small library CRISPR–SpCas9

We generated a small library targeting 369 genes to test in a mini-pool *in vivo* CRISPR/Cas9 screen. We prioritized: 1) proteomics hits with FDA available drugs and hits from the CMap analysis, 2) most statistically significant CRISPR depleted/enriched hits, 3) genes that had been previously related to DHODH in published in vitro studies, and 4) a panel of immune-related hits. Four guide RNA sequences were designed for each gene based on the Brunello library and 528 non-targeting control guide RNA sequences were included. After adding uniform adaptor sequences to each guide RNA, the oligo pool was synthesized via Twist Bioscience. 10 ng of synthesized oligo pool was amplified using Q5 polymerase then purified using the Qiagen PCR Purification Kit (#28104). 500 ng purified oligo were mixed with 5 μg lentiCRISPR v2 (Addgene #52961) backbone, which was linearized with Esp3I enzyme (NEB #R0734L). The ligation was performed by adding an equal volume of NEB HiFi DNA Assembly Kit (#E5520S) into the above mix and incubated at 50 °C for 60 min. Electrocompetent transformation was applied to the assembled products using ElectroMAX Stbl4 Competent Cells (Thermo #11635018) according to the standard manufacturer’s protocol. Transformed cells were incubated in 1 L Terrific Broth at 225 RPM 30 C for 14 h. ZymoPURE II Plasmid Maxiprep Kit (ZYMO #D4203) was used to extract plasmid DNA. The distribution degree of all guides in amplified library were assessed via NGS to ensure the Gini index is less than 0.1. Lentivirus production and infection was performed as described in the in vitro genome-wide screen. Virus infection ratio was controlled at 10%. For cell culture-based screens, 1000x depth of each guide was used. BQ was applied using the same approach as described in *in vitro* genome-wide screen. Genomic DNA was extracted at the end of treatment. For the *in vivo* screen, 5 million PaTu-8988T cells in 50 μl PBS and 50 μl matrigel were injected in bilateral flanks of NOG (NOD.Cg-*Prkdc^scid^ Il2rg^tm1Sug^*/JicTac) mice (n= 10 tumors per arm). This strategy ensured 4000x depth for each guide. Treatment was initiated when tumors reached 80-100 mm^3^. BQ was prepared as previously described^3^ by dissolving in 70% PBS 1x /30% PEG-400 and adjusting pH to 7.00 with NaOH. Mice were injected once every other day for 14 days at 50 mg/kg with BQ or vehicle. At endpoint (4h after last dose), tumor tissues were collected, and genomic DNA was extracted using DNeasy Blood & Tissue Kit (Qiagen #69504). NGS library preparation, sequencing, and analysis were performed as described for the genome-wide screen.

### Real time Cell Proliferation and Apoptosis

Annexin V conjugated to a fluoroprobe have been optimized for detection of phosphatidylserine (PS) externalization. Addition of the Incucyte® Annexin V Dyes to healthy cells yields little or no intrinsic fluorescent signal. Once cells undergo apoptosis, plasma membrane PS asymmetry is lost leading to exposure of PS to the extracellular surface and binding of the Incucyte® Annexin V Dye, making a bright and photostable fluorescent signal. The detection of cell growth and Annexin was performed following Incucyte®S3 protocols. Cells were plated at 1,000-2,000 cells/well in 96-well plates. The next day, drugs were added at the indicated concentrations. For apoptosis monitoring, Incucyte® Annexin V Red Dye was added at the same time as indicated drugs. Plates were kept in the Incucyte®S3 system inside a cell culture incubator for 72 hours. Cell growth and the accumulation of Annexin V fluorescent signal were captured with the Incucyte® Live-Cell Analysis System every 3-4 hours. The Annexin fluorescent signal was analyzed by Incucyte® Live-Cell Analysis System and the Total Red Object Integrated Intensity (RCU x µm²/Image, RCU means Red Calibrated Unit) was used to plot the curve in graphpad. For the cell growth analysis, relative cell growth (%) normalized to day 0 (day0 =100) was calculated by Incucyte® Live-Cell Analysis System and used to plot the curve in graphpad.

### Synergy testing

On day 0, 300 cells in 20 μl of medium were seeded in 384 well plates by a microplate dispenser (Type 836, Thermo Scientific Combi). HPAC was seeded at 500 cells/well. On day 1, drugs were administered with a Tecan D300e drug dispenser (Tecan, Männedorf, Switzerland). Plates were kept in an incubator for 5 days. 20 μl of CellTiter-Glo was added by microplate dispenser and plates were incubated on a shaker at room temperature for 1 h. Luminescence was measured with a Beckman Coulter Cytoflex. IC50 values were calculated using non-linear regression analysis in Graphpad Prism. Synergy score between the two drugs was calculated with the viability (%) which normalized to DMSO control using the HSA model implemented in SynergyFinder (https://www.synergyfinder.org/)^66^. The mean HSA value of drug combination was used to plot the synergy score heatmap by graphpad.

### Apoptosis analysis by flow cytometry

Cells were plated at 50,000 cells/well (24-well plate). Cells were treated the next day with the indicated compounds for 72 h. Adherent and non-adherent cells in the medium were collected and stained with Annexin-FITC and propidium iodide for 15 min (BD biosciences 556547) as per the manufacturer protocol. Cells were placed on ice and analyzed using a Beckman Coulter Cytoflex. Quantification of dead cells after treatment included cells which were in early apoptosis (FITC Annexin V positive and PI negative) as well as late apoptosis (FITC Annexin V and PI positive).

### BH3 profiling

BH3 profiling was conducted via flow cytometry following established protocols^67^ Briefly, cultured cells were trypsinized, centrifuged at 500xg for 5 minutes, resuspended in mannitol experimental buffer (MEB; 10 mM HEPES (pH 7.5), 150 mM mannitol, 50 mM KCl, 0.02 mM EGTA, 0.02 mM EDTA, 0.1% BSA, and 5 mM succinate), and added to wells of prepared 96-well plates containing the indicated peptide conditions and 0.001% digitonin. The cells were then incubated for 60 min at 28 °C, followed by fixation for 15 minutes in 8% PFA. Fixation was neutralized using N2 buffer (containing 1.7 M tris base and 1.25 M glycine, pH 9.1), and the cells were subsequently stained overnight with DAPI and an Alexa Fluor 647-conjugated anti-cytochrome c antibody (Biolegend, 612310) at 4 °C. Finally, the stained cells were analyzed using an Attune NxT flow cytometer. The percentage of cytochrome c negative cells was determined for each peptide treatment condition.

### Western blotting

Cells were collected in ice cold PBS and lysed in RIPA buffer containing 1 × Halt Protease and Phosphatase Inhibitor (Thermo Scientific, 78446). Lysates were centrifuged and supernatants collected. Protein (100 μg) was resolved on 4% to 12% SDS-PAGE gels and transferred to PVDF membranes. Membranes were blocked in 5% milk and incubated with primary antibodies followed by peroxidase-conjugated secondary antibodies. Membranes were developed using the ECL Detection System (Thermo, 32209). The following antibodies were used: BCL-2 (CST, #3498, 1:500), BCL-X_L_ (CST, #2764, 1:1000), MCL-1 (CST, #5453S, 1:1000), PARP (CST, 9542S, 1:1000), PUMA (CST, #98672S, 1:1000), BIM (CTS, #2933, 1:1000), cleaved caspase 3 (CST, #9661, 1:1000), ACTB (Sigma, A5441, 1:5,000), Anti-rabbit IgG (H1L) HRP conjugate (Thermo, 31460, 1:3,000), and Anti-mouse IgG (H1L) HRP conjugate (Promega, W4021, 1:7,000). Quantification was performed using ImageJ.

### Subcutaneous Mouse Xenograft and Allograft Studies

Animal studies were performed in accordance with a Dana-Farber Cancer Institute Institutional Animal Care and Use Committee–approved protocol (10-055). Mice were housed in pathogen-free animal facilities at Dana-Farber Cancer Institute. PaTu-8902 cells (1 × 10^6^ in PBS:Matrigel (1:1)) were implanted subcutaneously in the flank of NOD.Cg-*Prkdc^scid^ Il2rg^tm1Sug^*/JicTac mice (NOG mice, 7 weeks, female, Taconic). Tumors were allowed to reach 150 mm^3^ before randomization to treatment groups. HPAC (3 × 10^6^ in PBS:Matrigel (1:1)) cells were implanted subcutaneously in the flank of the NOD.Cg-*Prkdc^scid^ Il2rg^tm1Wjl^*/SzJ mice (NSG mice, 8 weeks, female, Jackson Lab). Tumors were allowed to reach 150-200 mm^3^ before randomization to treatment groups. KPCY 6694 C2 cells (0.5 × 10^6^ in PBS:Matrigel (1:1)) were implanted subcutaneously in the flank of C57BL/6J mice (8 weeks, female, Jackson Lab). Tumors were allowed to reach 75 mm^3^ before randomization to treatment groups. DT2216 was formulated in 50% phosal 50 PG / 45% miglyol 810N / 5% polysorbate 80. BQ was formulated in 70% PBS 1x /30% PEG-400 and adjusting pH to 7.00 with NaOH. DT2216 (15 mg/kg, twice per week) and BQ (10 mg/kg, three times per week) were administered intraperitoneally for 21 days. Mice in vehicle-control arms received 100 μl 30% PEG-400 / 70% PBS and 100 μl 50% phosal 50 PG / 45% miglyol 810N / 5% polysorbate 80 following the same schedule as DT2216 and BQ administration. Tumor volume was measured twice a week with calipers as follows: volume = (length × width^2^)/2. Tumors were then removed and cut into several pieces which were frozen or fixed in 10% formalin for future analysis. Drug toxicity was assessed by weighing mice twice a week and by complete blood counts at endpoint. Blood was drawn by sub-mandibular bleeding and analyzed using GENESIS^TM^ hematology analyzer from Oxford Science.

### Orthotopic models of PDAC

Orthotopic surgeries were performed as described^68^. 50,000 KPCY 6694 C2 cells in 30 μl of PBS:Matrigel 1:1 were implanted in pancreata of C57BL/6J mice. DT2216 (15 mg/kg, twice per week) and BQ (10 mg/kg, three times per week) were administered intraperitoneally starting on day 4 after implantation. Mice were weighed twice a week to assess drug-related toxicity. Mice were euthanized 20 days post-surgery and tumors were harvested at endpoint and weighed.

### Immunohistochemical staining

Tissues were processed as previously described^15^. Tissues were fixed in formalin and paraffin embedded. After deparaffinization, primary antibody was incubated followed by secondary antibody and then developed by DAB. The following antibody was used: cleaved caspase-3 (Asp175) (CST, 9661,1:400). Positive cell staining was quantified by ImageJ. Four 20× fields per mouse were analyzed.

### Bioinformatic Analysis

Quantitative proteomics dataset analyses were performed as described^14^. Datasets were subject to analysis using LIMMA package (3.40.2)^69^ in R platform. Between each pairwise comparison, LIMMA applied a standard linear model fitting and an empirical Bayes procedure to correct the distribution. After the correction, a moderated t-statistic was calculated for each gene using a simple Bayesian model, which served as the fundamental statistic. Type I error is corrected by calculating false discovery rate (FDR) using the Benjamini-Hochberg method. A volcano plot of log_2_ fold change and FDR served as a visualization method for each comparison. Broad GSEA software (3.0)^70^ was used to perform gene set enrichment analysis (GSEA) against the collection containing all canonical pathways (c2.all.v2023.2.Hs.symbols.gmt) in MSigDB. Pathway clouds were visualized in Cytoscape (3.7.2)^71^ (with EnrichmentMap (3.2.1)^72^ plugin, and each cluster was manually curated. A target list and a background list were created for each comparison and then used for Gene Ontology (GO) enrichment analyses^73^. Principal component analysis (PCA) was done with the prcomp function in R platform, and the two major components (PC1 and PC2) were used for visualization. The L1000 gene expression database in Connectivity Map (CMap) 2.0^22^ was used as the reference and top 150 up- and downregulated proteins upon BQ treatments were queried against L1000 to calculate connectivity scores. To identify BCL-XL, MCL1 and BCL2 dependencies and also BCL-XL expression across 45 PDAC cell lines, we analyzed data from pooled, genome-scale CRISPR–SpCas9 loss-of-function data within the Broad Institute’s DepMap Public 23Q4+Score (Chronos) and RNAseq data within expression Public 23Q4^35^.

### Chemicals

The following chemicals were used: Brequinar (MCE, HY-108325), BAY-2402234 (MCE, HY-112645), A-1331852 (MCE, HY-19741), DT2216 (Chemietek), uridine (Sigma, U3003), Z-VAD-FMK (Selleckchem, No.S7023), FITC Annexin V Apoptosis Detection Kit I (BD Biosciences, 556547), Incucyte® Annexin V Dye (Sartorius, 4641), Normal goat serum (Vector Labs, S-1000), Vectastain ABC-HRP kit (Vector Labs, PK-4001), Goat Anti-Rabbit lgG(H+L), Biotinylated (Vector Labs, BP-9100), and ImmPACT DAB Substrate kit (Vector Labs, SK-4105), CellTiter-Glo® 2.0 Cell Viability Assay (Promega, G924).

### Statistical Analysis

For comparisons between two groups, Student t*-*test (unpaired, two-tailed) was performed as noted. Groups were considered different when *P* < 0.05. For multiple comparisons, ordinary one-way ANOVA tests (Tukey statistical hypothesis test was used for correlation) were performed using Prism 8 (GraphPad Software). R-based analyses as described above, and figure generation were performed in R v4.0.3.

### Illustrations and Diagrams

Drawings for experimental set-up were created in Adobe Illustrator (v24.1.2) and BioRender.

## Acknowledgments

This research was supported by a Burroughs Wellcome Fund Career Award for Medical Scientists and the Sidney Kimmel Foundation Kimmel Scholar Program (to JDM), a Claudia Adams Barr Program for Innovative Cancer Research grant (to NSC), and the Hale Family Center for Pancreatic Cancer Research (to JDM, SKD, AJA). We thank Harvard Medical School Rodent Histopathology core for tissue processing and histopathological interpretation services. We acknowledge Dr. Steven Gygi for use of CORE for mass spectrometry data analysis software.

## Data availability

LC–MS, MS and NGS data present in this study have been deposited as supplementary tables. The other data supporting the findings of this study are available within the article and its Supplementary Information files.

