## Supplementary Information for "De novo pyrimidine biosynthesis inhibition synergizes with BCL-X_L_ targeting in pancreatic cancer"

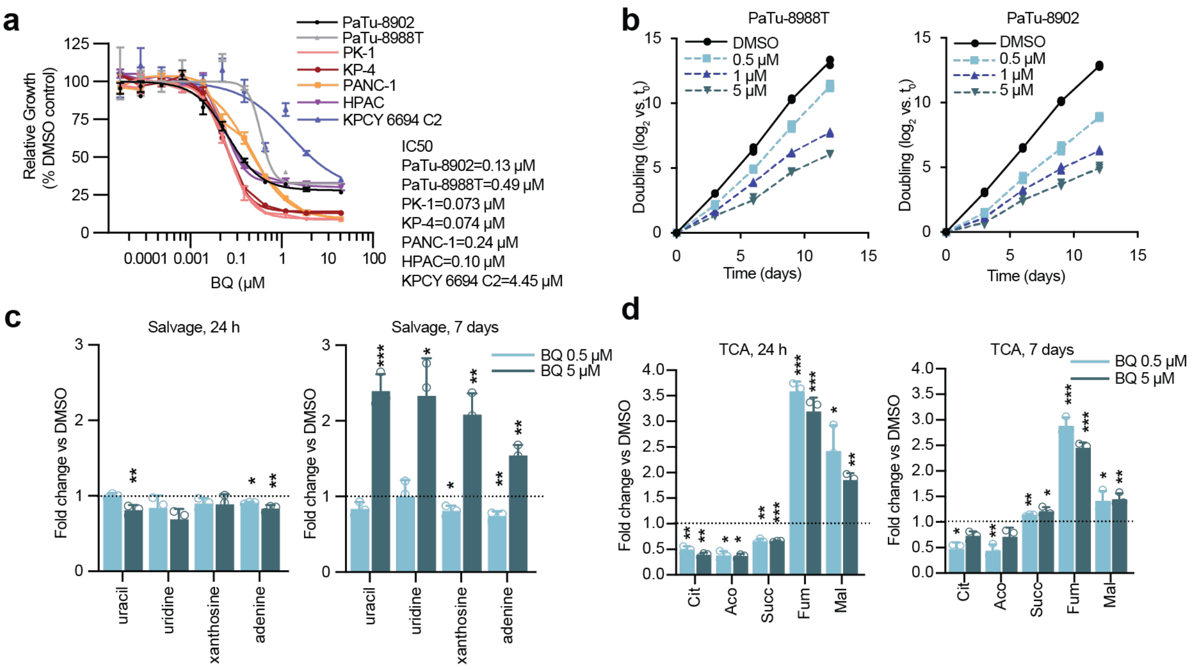


**Supplementary Figure 1 Metabolomics and proteomics identify compensatory mechanisms to BQ** **(a)** Cell proliferation dose-response curves for PDAC cell lines treated with BQ in DMEM. Error bars represent s.d. of two technical replicates. **(b)** Doubling curves for PaTu-8988T and PaTu-8902 cells treated with BQ (0, 0.5 μM, 1 μM, 5 μM). **(c)** Fold change of metabolites in the nucleotide salvage pathway correlate with a compensatory activation of the pathway at 7 days (24h: left panel, 7 days: right panel) in PaTu-8988T cells. **(d)** Fold change of metabolites in the Tricarboxylic acid (TCA) pathway for PaTu-8988T cells. Cit: citrate; Aco: aconitate; Succ: succinate; Fum: fumarate; Mal: malate.

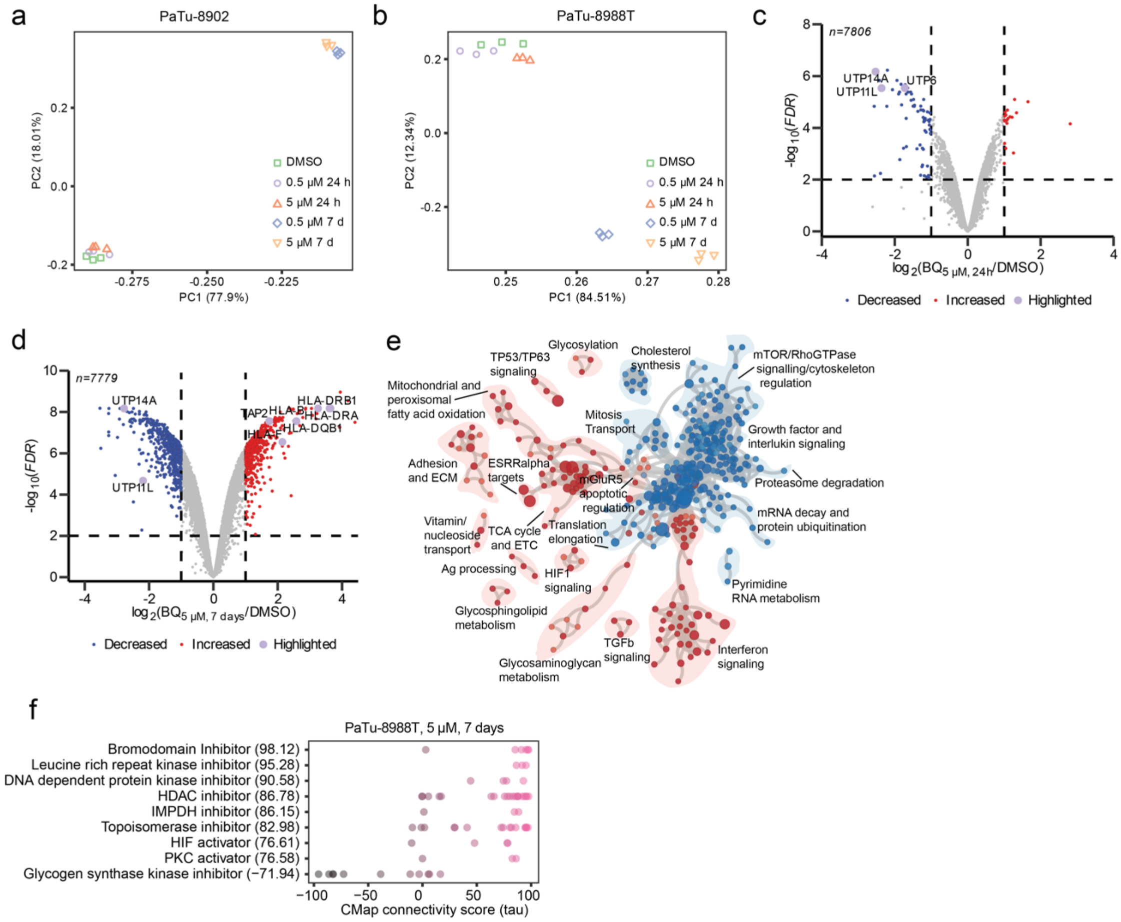


**Supplementary Figure 2 Quantitative temporal proteomics of BQ-treated PDAC cells**

**(a-b)** Principal Component Analysis plots of proteomics data for DMSO or BQ-treated PaTu-8902 **(a)** and PaTu-8988T **(b)** cells show sample clustering. **(c-d)** Volcano plot illustrates significant protein abundance differences in PaTu-8902 cells treat with BQ 5 μM at 24h **(c)** and 7 days **(d)**. Volcano plots display the -$\log_{10}$ (FDR) versus the $\log_2$ of the relative protein abundance of mean BQ to DMSO-treated samples. Purple circles highlight proteins previously linked to DHODH function. Red circles represent significantly upregulated proteins ($\log_2$ fold change≥1), whereas blue circles represent significantly downregulated proteins ($\log_2$ fold change≤-1; data from 3 DMSO or 3 BQ-treated independent plates). **(e)** Enrichment map of gene set enrichment analysis (GSEA) of BQ-proteomes from PaTu-8902 cells at 7 days (0.5 μM). FDR<0.01, Jaccard coefficient>0.25, node size is related to the number of components identified within a gene set and the width of the line is proportional to the overlap between related gene sets. GSEA terms associated with upregulated (red) and downregulated (blue) proteins are colored accordingly and grouped into nodes with associated terms. **(f)** Connectivity map analysis for PaTu-8988T cells with BQ (5 μM, 7 days). Perturbagen classes with mean connectivity scores >90% or <-90% and FDR<0.05 are displayed.

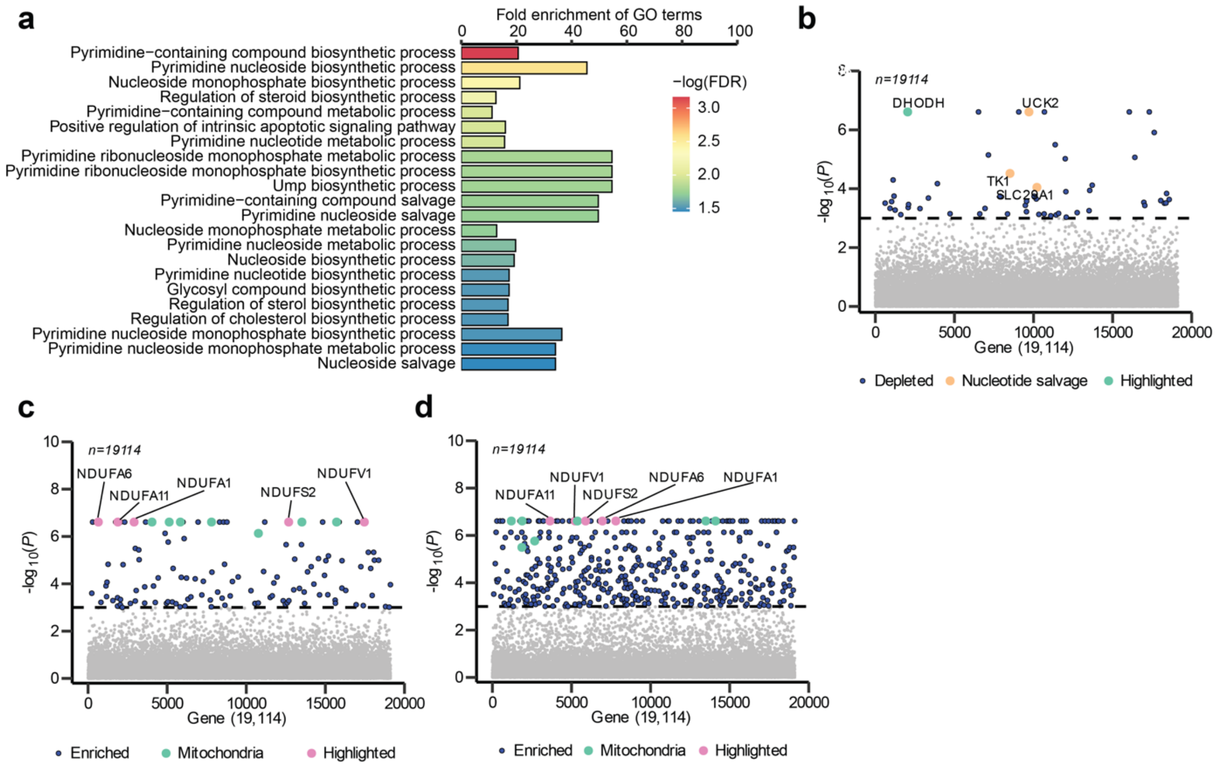


**Supplementary Figure 3 BQ-anchored genome-wide CRISPR/Cas9 screen identifies combinatorial drug targets in PDAC cells**

**(a)** Gene Ontology analysis of the normalized top 100 depleted genes from the in vitro CRISPR/Cas9 screen in PaTu-8988T cells (BQ 5 μM) classified by p-value. **(b-d)** Manhattan plot for in vitro CRISPR screen in PaTu-8988T cells shows **(b)** depleted hits at 0.5 μM BQ, and **(c)** enriched hits at 0.5 μM or **(d)** 5 μM BQ**.** For all plots, blue dots highlight significant genes ($-\log_{10}(P)=3$) in BQ versus DMSO.

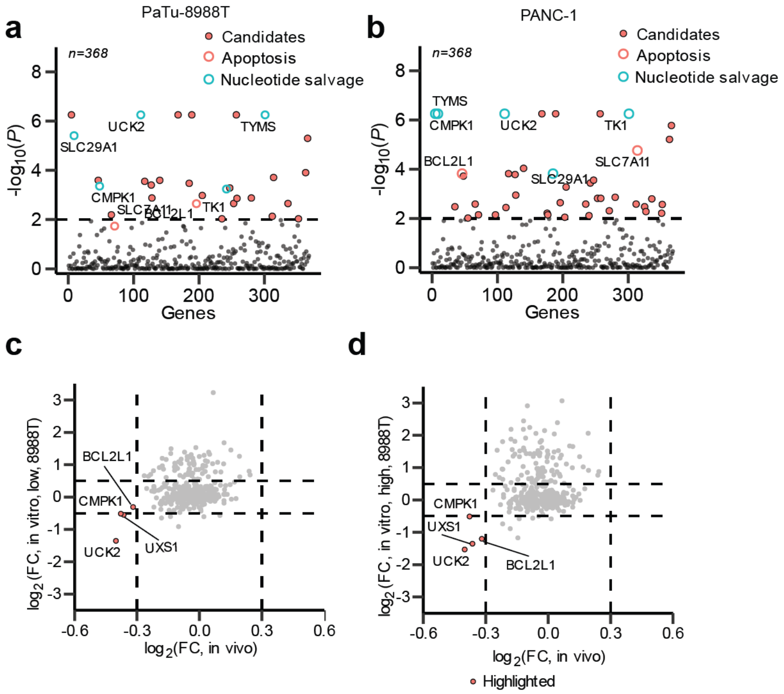


**Supplementary Figure 4 BQ-anchored genome-wide loss-of-function CRISPR/Cas9 screen identifies combinatorial drug targets in PDAC cells lines**

**(a-b)** Manhattan plot for depleted hits in a mini-library in vitro CRISPR screen in PaTu-8988T **(a)** and PANC-1 **(b)** cells at 0.5 μM BQ. **(c-d)** Correlation plot of depleted or enriched hits in in vitro whole-genome CRISPR/Cas9 screen vs. in vivo depleted hits in PaTu-8988T tumors at low or high BQ doses.

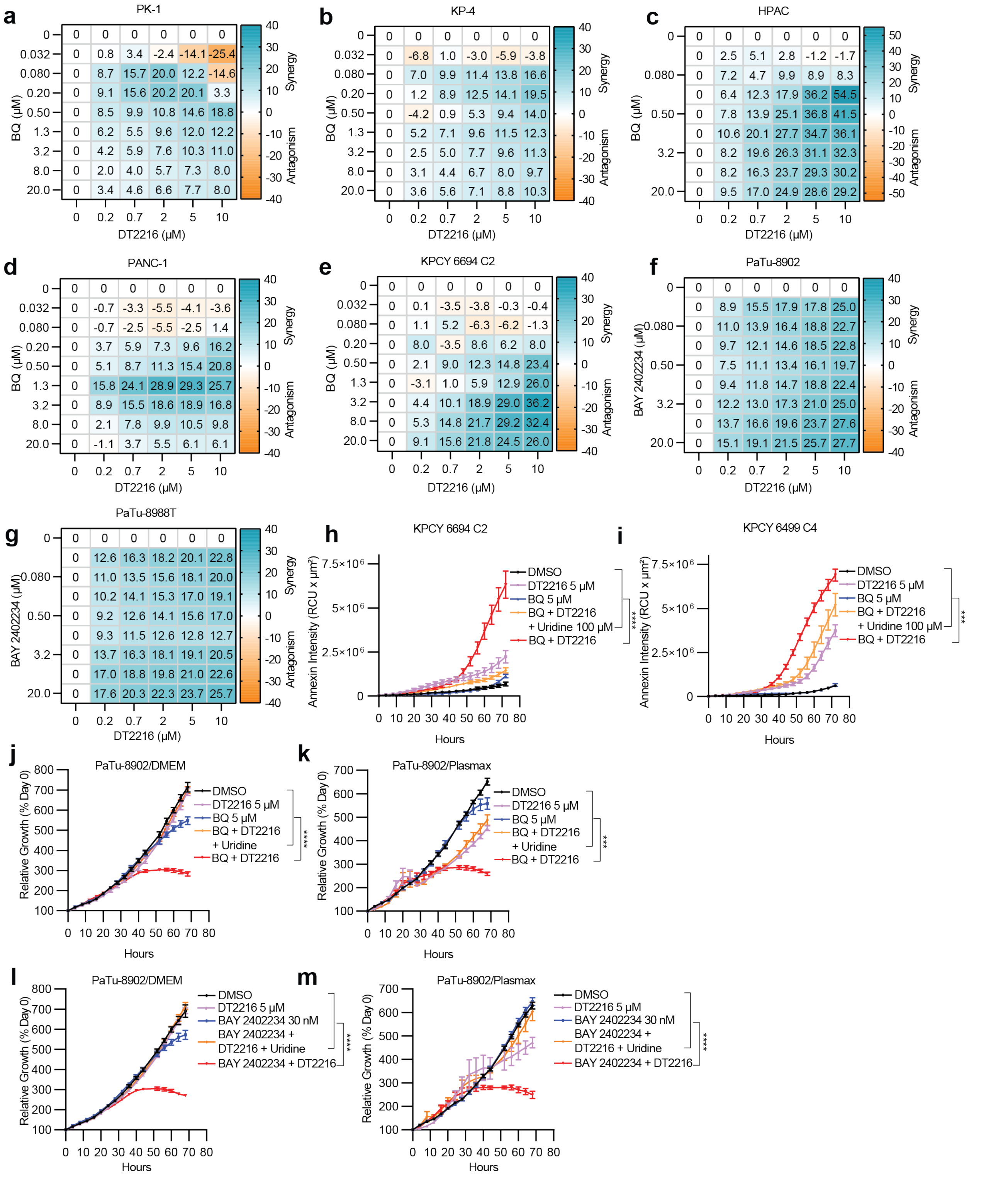


**Supplementary Figure 5 BQ and DT2216 combination demonstrates synergy in PDAC cell lines**

**(a-g)** Synergy score heatmaps of combination treatment with DT2216 and BQ (a-d) or DT2216 and BAY2402234 (f-g) in PDAC cells. Synergy score between the two drugs was calculated using the HSA model implemented in SynergyFinder (antagonism: <− 10; additive effect: from − 10 to 10; synergistic effect: >10). Data shown as mean HSA score of three technical replicates. **(h-i)** Real-time Annexin signal accumulation in KPCY 6694 C2 and KPCY 6499 C4. Cells were labeled with Annexin V Red Dye and treated with the indicated concentrations of BQ or DT2216 (alone or in combination) or in combination with uridine (100 μM) for 72 hours. **(j-k)** Relative proliferation of PaTu-8902 cells in different media (DMEM **j**, Plasmax **k**) treated with the indicated concentrations of BQ or DT2216 alone or BQ + DT2216 or BQ + DT2216 in combination with uridine (100 μM) for the indicated time points. **(l-m)** Relative proliferation of PaTu-8902 cells in different media (DMEM **l**, Plasmax **m**) treated with the indicated concentrations of BAY 2402234 or DT2216 alone or BAY 2402234 and DT2216 or BAY 2402234 and DT2216 in combination with uridine (100 μM) for the indicated time points. Real-time cell growth was monitored by incucyte. Error bars represent s.d. of three technical replicates (representative of two independent experiments). For all panels, significance determined with ordinary one-way ANOVA. *$p < 0.05$, **$p < 0.01$, ***$p < 0.001$, ****$p < 0.001$.

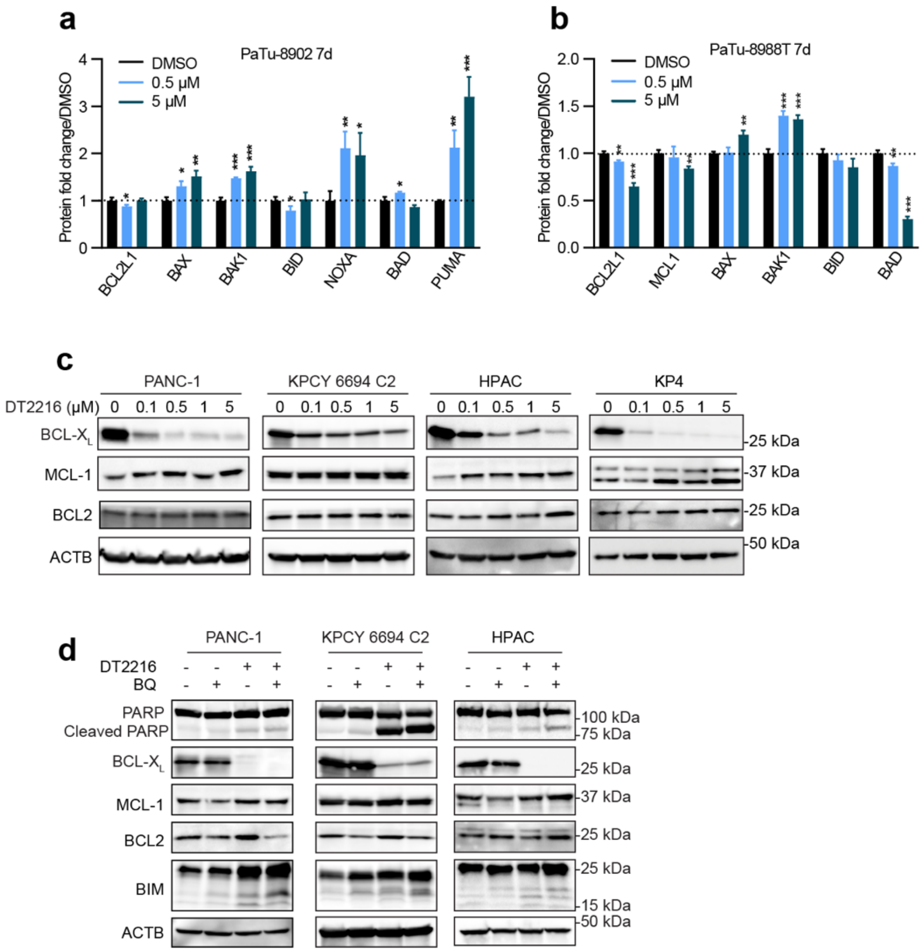


**Supplementary Figure 6 Protein level changes of BCL-2 family proteins with BQ or DT2216 alone or BQ and DT2216 combination treatment**

**(a-b)** Proteomics data derived from Fig. 1e and Supplementary Fig. 2d of BCL-2 family protein levels in PaTu-8902 and PaTu-8988T cells treated with 5 μM BQ for 7 days (fold change vs. DMSO). Statistical significance was determined by Student t-test. $^{*}p < 0.05$, $^{**}p < 0.01$, $^{***}p < 0.001$. **(c)** Immunoblot analysis of BCL-$X_L$, MCL1 and BCL2 in lysates from PANC-1, KPCY 6694 C2, HPAC, and KP4 cells treated with DMSO or DT2216 with indicated doses for 16 hours. **(d)** Immunoblot analysis of BCL-2 family proteins in lysates from HPAC, PANC-1 and KPCY 6694 c2 cells treated with DMSO or with DT2216 (5 μM) or BQ (5 μM) alone or in combination for 24 hours.

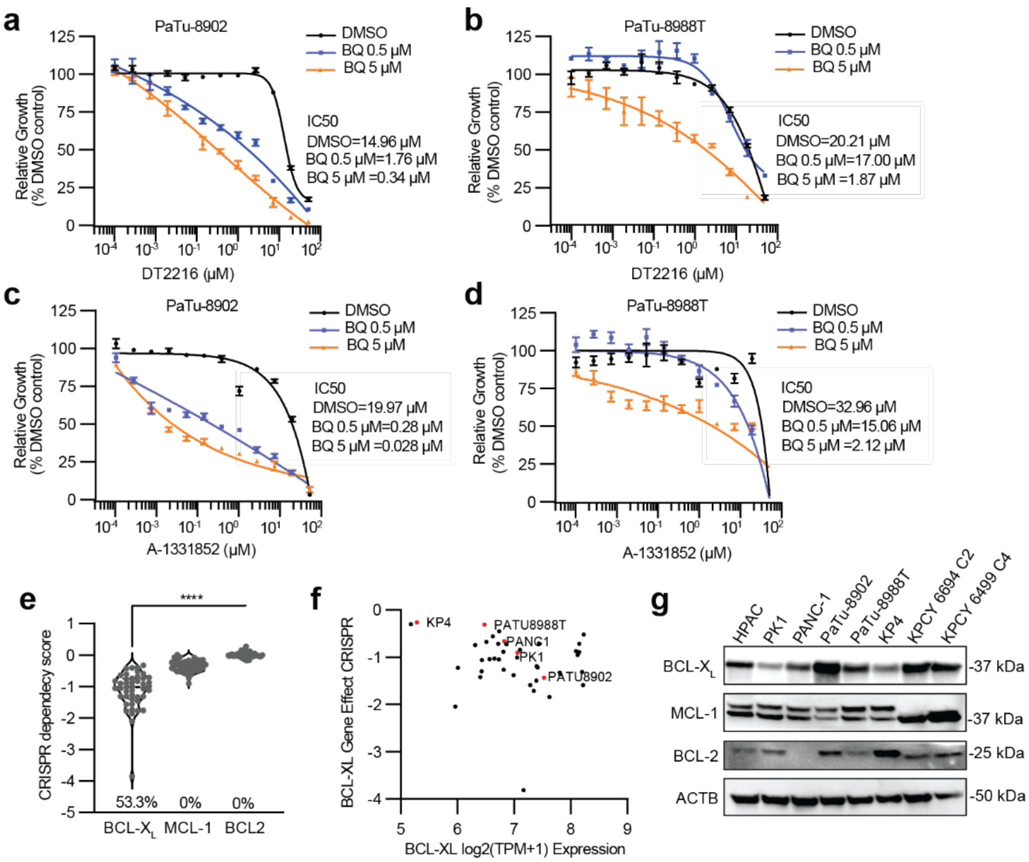


**Supplementary Figure 7 BCL-X$_L$ is a dependency in PDAC cell lines and BQ selectively increases sensitivity to BCL-X$_L$ inhibitors in PDAC cell lines**

**(a-b)** Percentage viability of PaTu-8902 and PaTu-8988T cells after they were treated with increasing concentrations of DT2216 with BQ 0.5 μM or 5 μM for 5 days. IC50 values are shown for a representative experiment out of three independent experiments. **(c-d)** Percentage viability of PaTu-8902 and PaTu-8988T cells treated with increasing concentrations of A-1331852 in combination with BQ 0.5 μM or 5 μM for 5 days. IC50 values are shown for a representative experiment out of three independent experiments. **(e)** Plot of CRISPR dependency scores of *BCL2*, *MCL1* and *BCL-X$_L$* in PDAC cell lines in the Cancer Dependency Map (DepMap) ($n = 45$); dashed line: median value; dotted line: quartile values. Percentage of cell lines scored as dependent (Dependency score less than −1) indicated at bottom of graph ($P < 0.0001$ for all comparisons to *BCL-X$_L$* using one-way ANOVA test)**.** **(f)** Plot of CRISPR dependency scores and mRNA expression level of *BCL-X$_L$* of PDAC cell lines in the Cancer Dependency Map (n=43); red dots indicate the cell lines evaluated in this study. **(g)** Immunoblot analysis of BCL-X$_L$, BCL2, and MCL1 in lysates from different PDAC cells, as indicated.

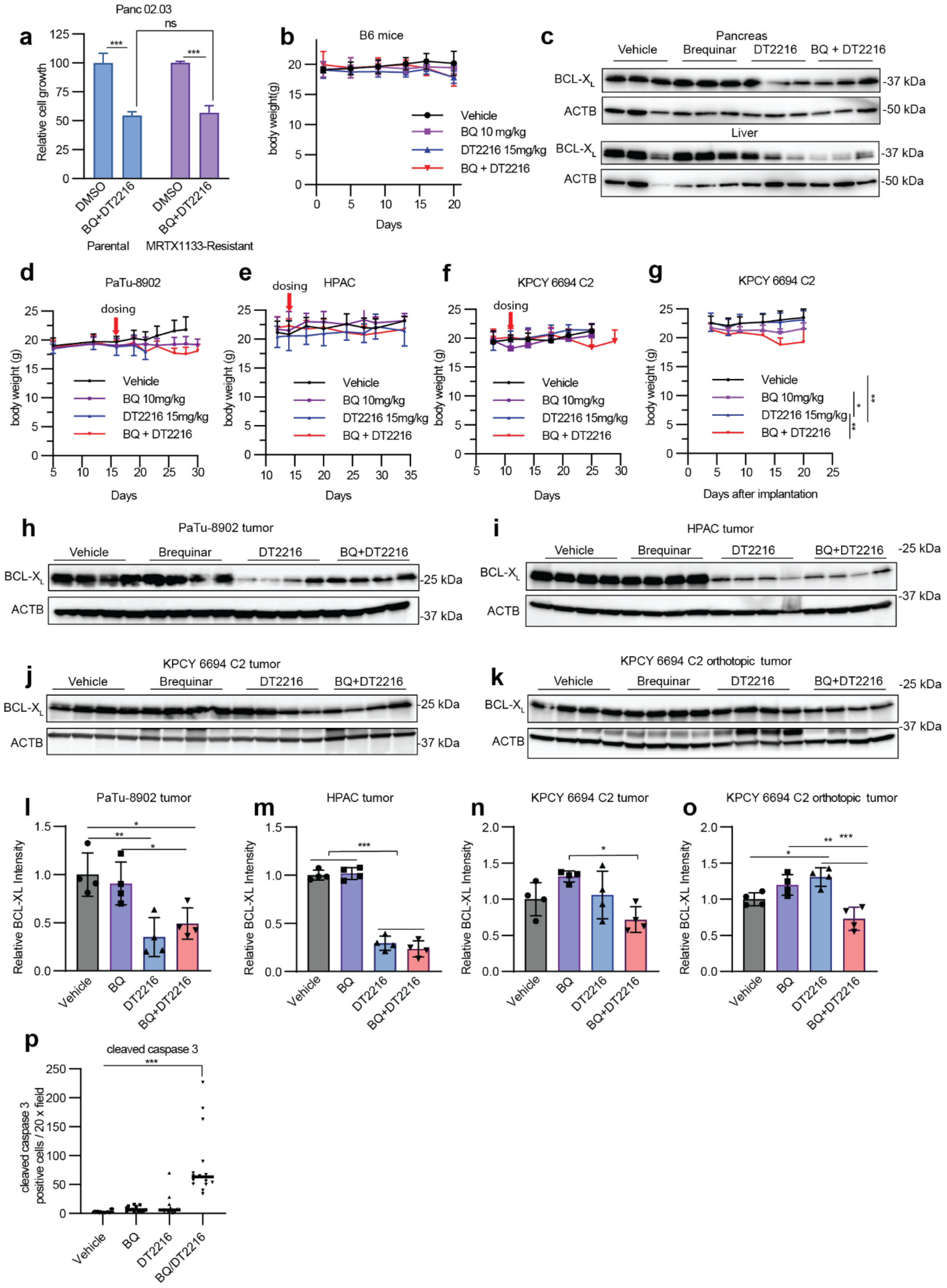

a
Panc 02.03
Relative cell growth
DMSO
BQ+DT2216
Parental
MRTX1133-Resistant
ns
b
B6 mice
body weight(g)
Days
Vehicle
BQ 10 mg/kg
DT2216 15mg/kg
BQ + DT2216
c
Pancreas
Vehicle
Brequinar
DT2216
BQ + DT2216
BCL-XL
ACTB
Liver
-37 kDa
-50 kDa
d
PaTu-8902
dosing
e
HPAC
f
KPCY 6694 C2
g
KPCY 6694 C2
Days after implantation
h
PaTu-8902 tumor
i
HPAC tumor
j
KPCY 6694 C2 tumor
k
KPCY 6694 C2 orthotopic tumor
-25 kDa
l
PaTu-8902 tumor
Relative BCL-XL Intensity
Vehicle
BQ
DT2216
BQ+DT2216
m
HPAC tumor
n
KPCY 6694 C2 tumor
o
KPCY 6694 C2 orthotopic tumor
p
cleaved caspase 3
cleaved caspase 3 positive cells / 20 x field
BQ/DT2216

**Supplementary Figure 8 Pharmacodynamic and tolerability evaluation of BQ and DT2216 combination therapy in vivo**

**(a)** Relative growth of Panc 02.03 and Panc 02.03 MRTX1133 resistant cells treated with vehicle (DMSO) or BQ and DT2216 for 3 days. Statistical significance was determined by Student t-test. ***p < 0.001. **(b)** Body weight changes of non-tumor bearing C57Bl/6J mice treated with vehicle, DT2216, BQ, or BQ and DT2216 for three weeks. **(c)** Immunoblot analysis of BCL-$X_L$ in pancreas and liver from non-tumor bearing C57Bl/6J mice treated with vehicle, DT2216, BQ, or BQ and DT2216 in combination as in **(b). (d-g)** Body weight of tumor bearing models treated as indicated and as presented in Figure 6: **(d)** PaTu-8902 flank xenograft, **(e)** HPAC flank xenograft, **(f)** KPCY 6694 C2 flank syngeneic allograft, **(g)** KPCY 6694 C2 orthotopic syngeneic allograft. **(h-k)** Immunoblot analysis of BCL-$X_L$ levels in xenograft and syngeneic tumors treated with vehicle, DT2216, BQ, or BQ and DT2216 in combination with indicated doses for 3 weeks. **(i-o)** Quantification of immunoblots in **(h-k). (p)** Number of cleaved caspase 3 positive cells was quantified in 4 fields (20X) from each tumor (PaTu-8902 tumor, *n* = 4 tumors per group). Statistical significance **(g, i-o)** was determined by one-way ANOVA. *p < 0.05, **p < 0.01, ***p < 0.001.
